# Motor Cortical Computations Underlying Natural Dexterous Movement in Freely Flying Bats

**DOI:** 10.64898/2026.03.25.714017

**Authors:** Boaz Styr, Kevin K. Qi, Xing Chen, William A. Liberti, Michael M. Yartsev

## Abstract

Elucidating the neural computations underlying natural, complex movement remains a fundamental challenge in neuroscience. Bat flight presents a formidable motor control challenge, requiring the use of hand-like wings whose many degrees of freedom must be precisely coordinated to enable rapid three-dimensional maneuvers. Here we performed large-scale wireless recordings of neuronal ensembles from the wing motor cortex of freely flying bats using Neuropixels probes, alongside detailed 3D pose tracking of wing kinematics. Despite the complexity of flight control, bats repeatedly executed highly accurate flights through precise adjustments of individual wingbeats. Surprisingly, motor cortical activity was not dominated by the global wingbeat cycle. Instead, individual neurons were sparsely active, exhibiting mixed selectivity for specific flight kinematics combined with variable entrainment to the wingbeat phase reaching millisecond-scale precision. This yielded a high-dimensional population regime driven by low shared variance across wingbeats, with successive wingbeats occupying distinct neural population states. Our findings reveal that during complex natural behavior the mammalian motor cortex operates in a high-dimensional computational regime that challenges prevailing views of motor cortical computation and underscores the importance of studying ethologically relevant behaviors to uncover neural principles governing brain function.

## Introduction

Uncovering the neural computations that govern natural, complex movements is a central goal of neuroscience. Studies in humans and animal models have established the central role of motor cortex in controlling complex movements^1–8^. Yet, much of what we know about how the mammalian motor cortex supports these behaviors comes from subjects trained on tasks that intentionally restrict behavioral degrees of freedom to produce movements that are amenable to quantitative analysis^9–16^. While such paradigms are invaluable, they may not fully engage the neural computations required for natural dexterous control. Indeed, studies using these approaches have consistently reported neural dynamics that appear to reside in a low-dimensional space that is largely invariant to the fine details of movement. In contrast, real-world motor behavior is often high-dimensional and complex. This creates a paradox: why are the very aspects of fine movement control that depend on motor cortex only weakly reflected in its neural dynamics? More broadly, can the neural principles derived from constrained behaviors generalize to natural dexterous motor actions that brains evolved to perform?^17^ Thus, fully probing the computational landscape of the mammalian motor cortex requires behavioral paradigms that (i) engage complex motor control, (ii) involve natural, ethologically relevant actions, (iii) are reproducible, and (iv) are compatible with large-scale neural recordings.

Bats are the only mammals capable of self-powered flight^18^, a form of locomotion that imposes exceptional demands on motor control, requiring complex 3D maneuvering that must be performed with high precision at high speeds that can exceed 40km/h^19,20^. To achieve this, bats developed several key adaptations. Chief among these is the evolution of complex wing structures composed of elongated, flexible fingers, permitting control over approximately twenty joint angles in each wing, giving the order of bats its name - *Chiroptera* or “hand-wing”^21^ (Figure 1A). In practice, bats utilize the many degrees of freedom offered by their hand-like wings for agile control of flight, exhibiting high levels of dexterity and maneuverability^22–25^. Consistent with the complexity of flight control, recent mapping of the sensorimotor cortex in Egyptian fruit bats (*Rousettus aegyptiacus*) has revealed an enlarged representation of the wings^26^. Thus, bat flight is a natural and ethologically relevant movement that also offers a formidable computational challenge to execute accurately.

**Figure 1.**
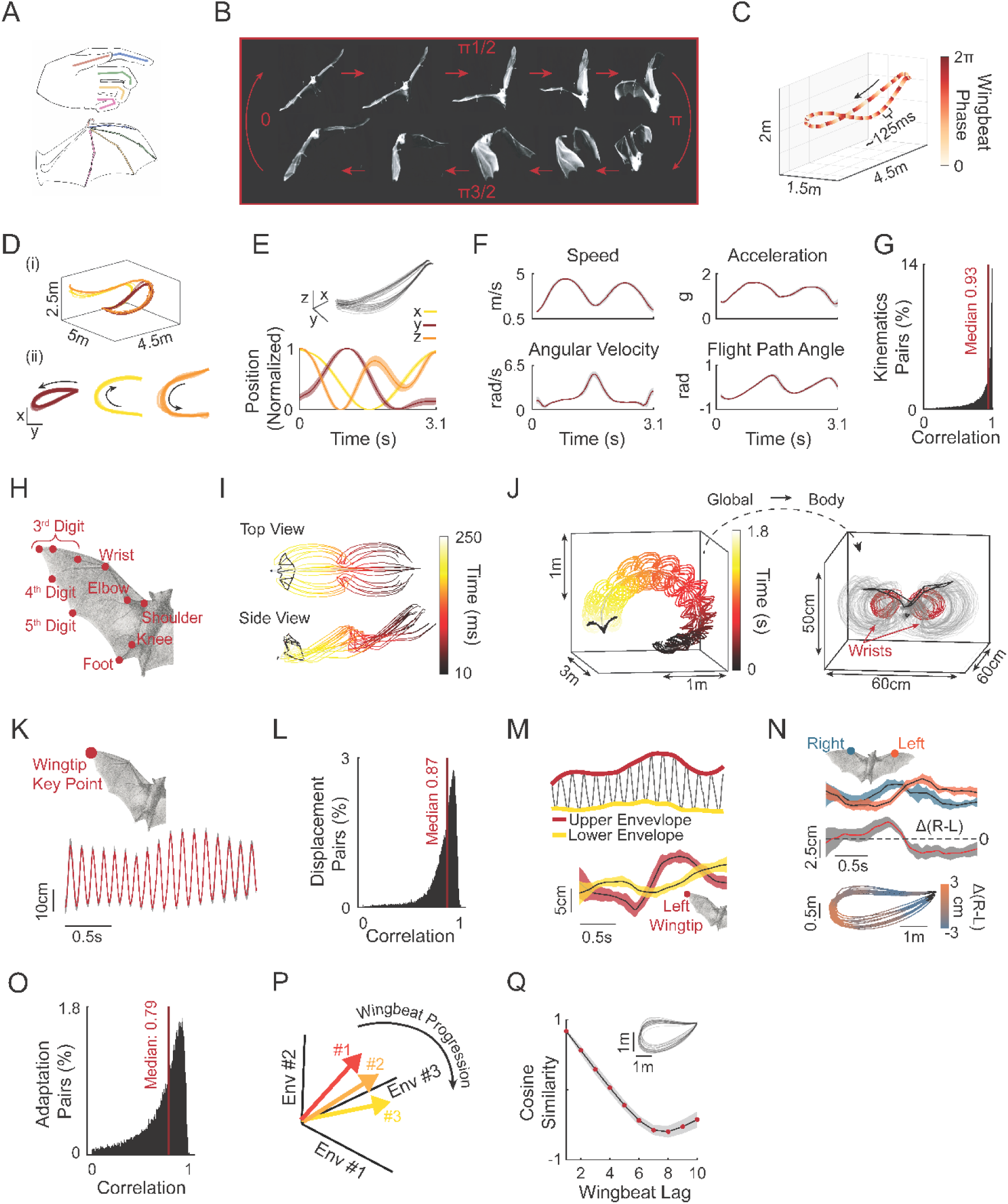
Flight reproducibility through precise wingbeat-level control. **A)** Schematic illustrating morphological homology between the human hand and the bat wing. **B)** Example frames from one tracking camera spanning a full wingbeat cycle. **C)** Example 3D position trace during a single flight. Black arrow indicates flight direction. Color indicates wingbeat phase. Wingbeats cycle ∼8 Hz (∼125 ms per cycle). **D)** Clustering flights into distinct paths. (i) 3D view of all clustered flights from one session. Colors represent distinct paths. (ii) Top-down projections of each of the three clustered paths. Black arrow indicates movement direction. Paths rescaled for visual clarity. **E)** Positional reproducibility for one example flight path. Top: all flights in 3D. bottom: x,y and z positions (range normalized 0-1). Shaded area represents SD. F) Four derived kinematic features from the same path depicted in (E) (speed, g-force, angular velocity and flight path angle). Gray shaded area represents SD. **G)** Distribution of correlations for position-derived kinematic features across flights of the same path (median 0.93; IQR; 0.20, n = 1,070 flights. p = 23 paths, N = 5 bats, F = 14 kinematic features). **H)** Illustration of key points used to train DeepLabCut model for 3D pose estimation. Depiction shows only the right side of bat body (10 of the 20 total key points used; Methods). **I)** Example of two spatial projections of the 3D pose estimation for two wingbeat cycles (x-y and x-z axis corresponding to top and side views, respectively). **J)** Example pose estimation of one flight in the global reference frame (left) and re-projected into the body reference frame (right). Two key points (left and right wrists) are shown in red. All other key points are in gray. **K)** Mean ± SD displacement across 32 flights of the same path for the right wingtip key point (3rd digit 3rd joint) in the body refence frame (projected to the dorsal ventral axis). Note the extremely low variation in key point positioning across all 32 flights. **L)** Distribution of correlations of displacements across all key points and paths (median corr.: 0.87; IQR: 0.15; p = 17 paths; f = 276 flights; N = 3 bats). **M)** Quantification of envelopes around displacements showing adaptations on top of the wingbeat cycle. Top: Example upper (red) and lower (yellow) envelopes. Bottom: mean ± SD envelopes for the wingtip across all flights of a single path. **N)** Right-left asymmetry between key points on the wings. Top: Mean ± SD of upper envelopes of the two wrist key points (blue- right wrist, orange – left wrist) showing distinct temporal evolutions along the same path. Middle: mean ± SD of the difference between the right and left wrists depicted above. Bottom: The same right-left difference projected onto the bat’s position (top-down view) showing shifts around the turn. **O)** Distribution of correlations of all envelopes and right-left asymmetry across all key points and paths (median correlation: 0.79; IQR: 0.31; p = 17 paths; f = 276 flights; N = 3 bats). **P)** Schematic showing wingbeat vectors in “wingbeat adaptation space”. Each wingbeat is represented by a vector of the mean values of envelopes and asymmetries during that cycle. Depiction showing three wingbeat vectors smoothly changing during flight. **Q)** Example of cosine similarity of the adaptation vectors as a function of wingbeat lag from a given wingbeat (path depicted in inset). Red dots represent mean for each wingbeat lag across flights. Black line and gray shaded area represent mean ± SD. across flights in the path.

To track flight at the level of the wingbeat, we established a multi-camera automated 3D markerless pose estimation system using deep learning based neural networks^27^. This allowed us to track bats with high temporal and spatial resolution while they flew freely inside a large flight room (Figure S1). In parallel, we wirelessly recorded neural activity from the bat wing motor cortex (Figure S2 and ref.^26^) using up to three Neuropixels probes simultaneously^28^, allowing us to record the activity of hundreds of well-isolated single units during spontaneous flight behavior. Together, this enabled measurement of motor cortical activity from single units to ensemble dynamics, alongside movement kinematics, during a highly ethological, unconstrained and complex motor behavior in a freely flying mammal.

## Results

### Flight reproducibility through precise wingbeat-level control

Bats spontaneously performed self-selected and self-paced flights. Each flight can be described as occurring across two nested timescales: (i) individual wingbeats consisting of fast bilaterally synchronous up- and down-stroke cycles occurring at approximately 8 Hz, which are then (ii) strung together in series to form complete flight paths typically reaching speeds of ∼4-5 m/s (Figure 1B-C). All bats (N = 5) spontaneously developed highly reproducible flight patterns that could be separated into distinct flight clusters, which we termed “paths”^29^ (Figure 1D, Methods). Since the same path can be transversed with varying kinematic (e.g., walking or running on the same route), we first asked how conserved the flight kinematics are that give rise to the observed spatial path reproducibility. We began by analyzing position-derived flight kinematics and found that across all bats and paths, flights required continuously modulating speed, acceleration, angular velocity, and flight path angle (Figure 1E-F; see additional examples in Figure S3). Despite the continuously varying kinematics of each individual flight, repeated flights of the same path were reproduced with extremely high fidelity (Figure 1F-G, median correlation 0.93; IQR; 0.20, n = 1,070 flights, p = 23 paths, N = 5 bats).

We next asked whether the observed flight kinematic precision requires equally tight control at the single wingbeat level. Optimal feedback control predicts that the motor system should only constrain dimensions that affect task goals, permitting variability elsewhere^30^. The degree to which wingbeat variability is restricted within the solution space that produces accurate flight paths in bats is an open question. To test this, we first examined wingbeat variability using onboard accelerometer^28^. Acceleration profile aligned with striking consistency across 20+ wingbeat cycles across flights of the same path (Figure S4A-B). Segmenting individual wingbeats from this signal (Figure S4A, Methods) revealed that wingbeat period remained remarkably constant within and across bats (median across bats: 122.6 ms; IQR: 6.9 ms; n = 13,897 wingbeats. Figure S4C) even as flight kinematics varied between paths (Figure 1D-E and Figure S3). Indeed, period consistency translated into consistency in the locations of consecutive wingbeats along flights of the same path (Figure S4D). These results indicate that accurate control over position derived kinematics operates primarily through modulation of wingbeat shape rather than wingbeat duration.

We next trained a DeepLabCut model on a total of 20 keypoints on both the body and the left and right wings (Figure 1H-I, Figure S1 and Movie 1). Transforming these into body-centered coordinates (Figure 1J) revealed that keypoints oscillated with the wingbeat cycle in a reproducible manner across flights of the same path (Figure 1K-L; median corr.: 0.87; IQR: 0.15; p = 17 paths; f = 276 flights; N = 3 bats, Figure S4E). However, bats continuously adjusted the amplitude of these movements as flights unfolded. We thus quantified these adjustments by extracting upstroke and downstroke envelopes, as well as their left-right asymmetries (Figure S4F, Methods). These features provide a simplified readout of the complex body kinematics that determine flight kinematics^24,25^. For example, the envelopes and the left-right asymmetries were strongly modulated around turns (Figure 1M-N). Importantly, even these adaptations were consistent across flights of the same path (Figure 1M-O, median corr.: 0.79; IQR: 0.31; p = 17 paths; f = 276 flights; N = 3 bats). Envelopes evolved smoothly but distinctly across wingbeats during flight (Figure 1P-Q): similarity between adjacent wingbeats was high but decayed rapidly, reaching low values within just a few cycles (Figure 1Q, Figure S4G). Thus, each wingbeat receives smoothly evolving, yet highly precise, control signals tailored to current flight demands.

Next, we set out to probe the motor cortical computations that might underlie this fine, cycle-by-cycle, motor precision during flight. To do so, we wirelessly recorded neural activity from the motor cortex of freely flying bats using Neuropixels probes (targeting the wing motor cortex (Figure 2A - adapted from ref. 24; Figure S2)). Across five bats, we isolated 2,813 units, with 81-416 units recorded simultaneously per session (Supplementary Table 1), capturing both single cell responses as well as population dynamics.

**Figure 2.**
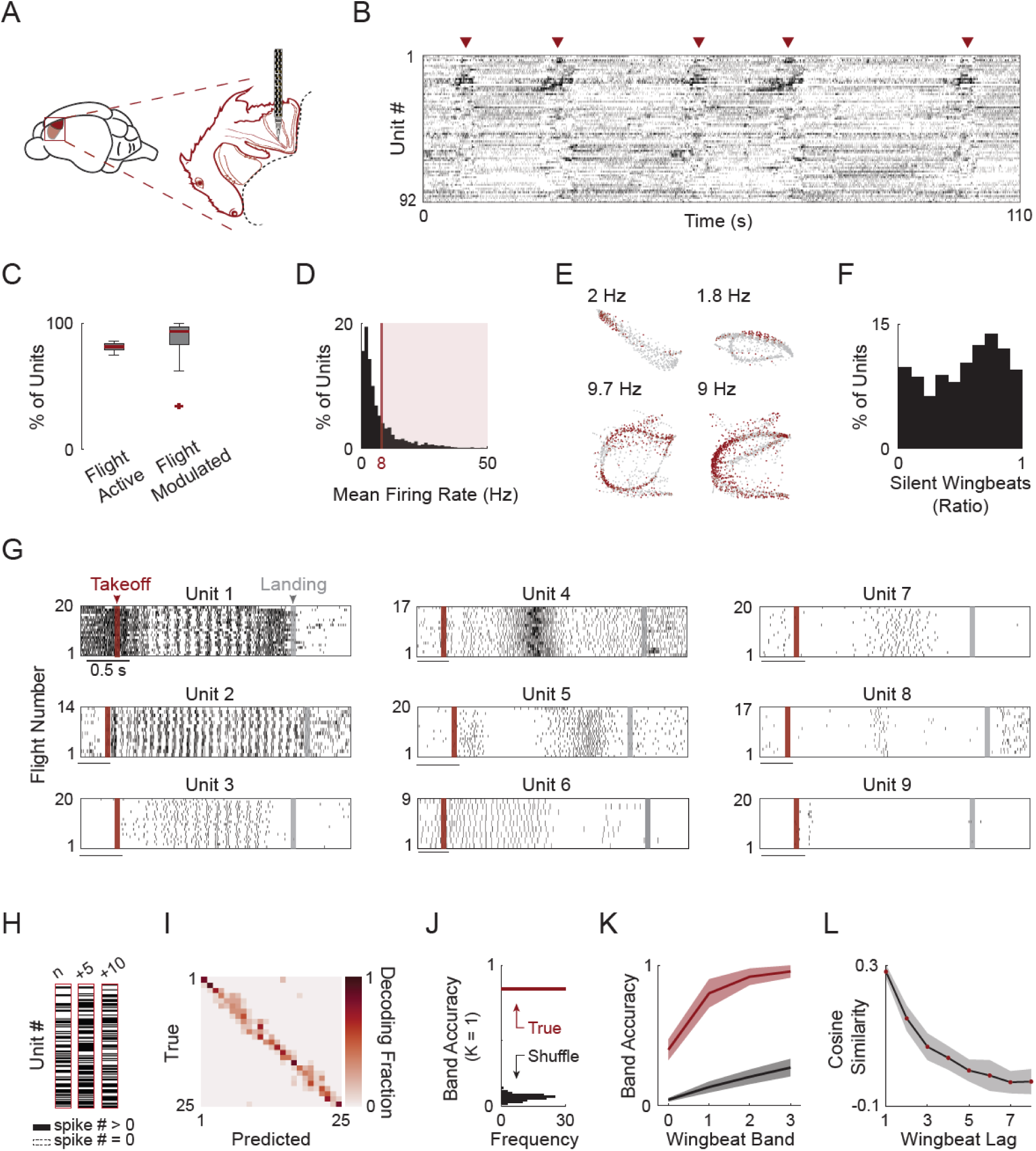
Selective recruitment of distinct neural subsets across wingbeats. **A)** Schematic showing the location of the motor cortex (light red) and wing motor area (dark red). Inset: Mapping of the bat motor cortex (“batunculus”) shows a large portion dedicated to the wing (modified from Halley et al. 2022). **B)** Raster plot of 92 simultaneously recorded units during a typical session. Spontaneous flights are indicated by red arrows. Units sorted using Rastermap. **C)** Fraction of flight active and flight modulated units (1,592/1,848 units modulated/flight active per session. S = 16 sessions. N = 5 bats). Box plots show median (red line), interquartile range (box), whiskers extending to 1.5× IQR, and outliers as individual red points. **D)** Distribution of mean firing rates during flight across all flight-modulated units. Shaded area indicates units with firing rates > 8 Hz. **E)** neural activity of four example units. Each dot represents the mean spatial position of the bat during a single wingbeat. Gray dots: no spikes; red dots: ≥ 1 spikes within the wingbeat. Spatial dimensions rescaled for clarity. **F)** Distribution of silent wingbeats (fraction of wingbeats without any spikes) across all flight modulated units. Higher silent wingbeat ratio translates to sparser units. **G)** Raster plots of nine example units across flights of a single path. Flights aligned to first wingbeat. Red line indicates first wingbeat, and gray line indicates mean time of last wingbeat corresponding to takeoff and landing, respectively. **H)** Examples of three wingbeat “barcodes” indicating which unit was active (≥ 1 spikes) for a given wingbeat. Numbers above barcodes indicate wingbeat number relative to the first. **I)** Confusion matrix from an SVM decoder trained to classify wingbeat groups within one example flight path using binary wingbeat vectors. Colors indicate mean 5-fold cross-validated decoding accuracy (n = 318 wingbeats, N = 25 wingbeat groups). **J)** Decoder mean accuracy (± 1 band around true group) in red, compared to the distribution of decoder accuracy for wingbeat group shuffled labels (100 iterations) in black. Same example path as in (I). **K)** Decoding accuracy as a function of band across all tested paths (n = 5 paths with > 100 units). Red: decoding using real group labels. Black: decoding from shuffled group labels. Lines and shaded areas indicate mean ± SD. **L)** Cosine similarity between population firing rate vectors as a function of wingbeat lag along the flight. Red dots: mean for each wingbeat lag across flights. Black line and gray shading: mean ± SD across flights in the path.

We considered two hypothetical regimes by which motor cortex might support complex natural dexterous movement (Figure S5). In one regime, motor cortical activity would largely follow the dominant structure of the behavior, tracking the wingbeat cycle, with single-neuron firing-rate modulations encoding adjustments superimposed on this rhythm. Population activity would therefore be broadly shared across cycles, yielding low-dimensional population dynamics (Figure S5A). This regime has been widely observed across constrained tasks^10,11,31–34^. Alternatively, neuronal activity might primarily reflect the adaptations occurring on top of the wingbeat rhythm. In this regime, motor cortex would selectively and sparsely recruit distinct neuronal subsets on individual wingbeats, such that a larger portion of neural variability reflects wingbeat cycle-specific activity. This would generate unique population patterns on successive wingbeats and, correspondingly, higher-dimensional population dynamics (Figure S5B). This regime maps more directly onto the established role of motor cortex in supporting dexterous movement^1,2,8,35,36^. We therefore set out to examine the different computational regimes that might characterize motor cortical activity, from the level of individual neurons to the population dynamics.

### Selective participation results in dynamic recruitment of distinct neural subsets across wingbeats

We first observed that neural activity in the motor cortex was distinctively modulated during flight (Figure 2B, sorted using Rastermap^37^). Starting at the level of individual flight active units (Methods) we initially assessed modulation by comparing the correlation of each unit’s activity across all flights in each path, to a null distribution obtained by circularly shifting the spikes times relative to the flight path independently. Most units significantly modulated their activity during flight (n = 1,592/1,848 or 86.1% of units modulated/flight active; Benjamini–Hochberg FDR corrected p-value, q = 0.01. Figure 2C) and many fired at rates well below the intrinsic ∼8 Hz wingbeat rhythm (Figure 2D) suggestive of a sparser activity regime than a smooth firing rate modulation aligned to the wingbeat cycle (Figure S5B). To further quantify this sparsity, for each unit, we computed the fraction of ‘silent wingbeats’, defined as the fraction of wingbeats during which no spikes were emitted. A typical session consisted of hundreds to thousands of individual wingbeats, providing extensive sampling of each unit’s activity across diverse kinematic conditions. We observed a great diversity of “on/off” patterns with some units only active on a small subset of wingbeats within a session (Figure 2E). Across all wingbeats and all sessions, about half of the units remained completely silent (no spikes) for half or more of all wingbeats (Figure 2F).

To further assess recruitment of units during the flight, we focused on activity patterns within specific paths. Aligning neural activity to flight onset revealed highly structured responses capturing both the complexity and the reproducibility of the flight behavior but with great diversity (Figure 2G). While some units exhibited clear phase-locking to the wingbeat cycle throughout the flight (e.g., units 1-3), others fired only during specific epochs of the flight at high precision (e.g., units 5-7). At the extreme, individual units fired just one or two spikes per flight yet repeated this ultra sparse, but highly precise pattern, across all flights of a given path (e.g., unit 9). These observations suggest that specific subsets of units might be recruited on specific wingbeats and at specific times (Figure S5A-B).

To systematically assess whether each wingbeat recruits a distinct population of units or whether the same units are shared across wingbeats, we clustered the wingbeats that occurred across flights on a given path by their kinematic similarity (Methods). This yielded groups that closely matched the number of cycles in a path and preserved their temporal order (Figure S6A). The advantage of this approach is that it captures even highly selective units active for just one cycle and allows averaging across flights by concatenating activity of individual wingbeat groups. This analysis revealed a more pronounced bimodal distribution of flight modulated units as compared to Figure 2F, with a clear peak for consistently active units and another peak for more selective ones (Figure S6B). We next asked whether selective recruitment would result in a unique activity pattern for different wingbeats along the flight. To directly test this, we constructed binary “barcodes” indicating which units were active (any spikes) versus silent for each wingbeat cycle (Figure 2H). Training a classifier to decode wingbeat group from these barcodes (Methods) yielded high accuracy, with misclassifications concentrated along the diagonal, suggesting that adjacent wingbeats recruit similar subsets of units and that the population of units which are active on each wingbeat evolves smoothly as flight unfolds (Figure 2 I-K). Directly quantifying barcode similarity confirmed this conclusion: Cosine similarity between population vectors decreased steadily, reaching stable values after several wingbeats (Figure 2L, Figure S6C). This rate of change matched the timescale at which wingbeat envelopes diverge (Figure 1Q, Figure S4G).

Together, these results indicate that motor cortex does not produce a fixed, continuously active population that modulates its firing rate on every wingbeat. Rather, it dynamically recruits distinct neuronal subsets on successive wingbeats in line with a mechanism of fine adaptations on top of a preexisting motor program (Figure S5). As consequence, the change in the neural subset that is recruited for each wingbeat mirrors the rate of modification on top of the wingbeat cycle and thus can potentially encode the adaptations required for each wingbeat.

### Motor cortical activity tracks kinematic demands across flight paths

We next reasoned that if indeed the neuronal population reflects fine adaptations for each wingbeat cycle, it should be sensitive to unique flight demands. To test this, we first asked whether different flight paths engage the same sequence of units. We leveraged the fact that bats naturally developed multiple distinct, yet highly reproducible, flight patterns. Comparing two paths of similar durations but opposite directions (Figure 3A) revealed path-specific neural activity. Sorting the simultaneously recorded units by their peak activity in one direction showed clear sequential structure, but this ordering failed to reveal the same sequential activity on the opposite direction (Figure 3A and B). Thus, each path recruits a unique neuronal sequence rather than activating a universal flight pattern.

**Figure 3.**
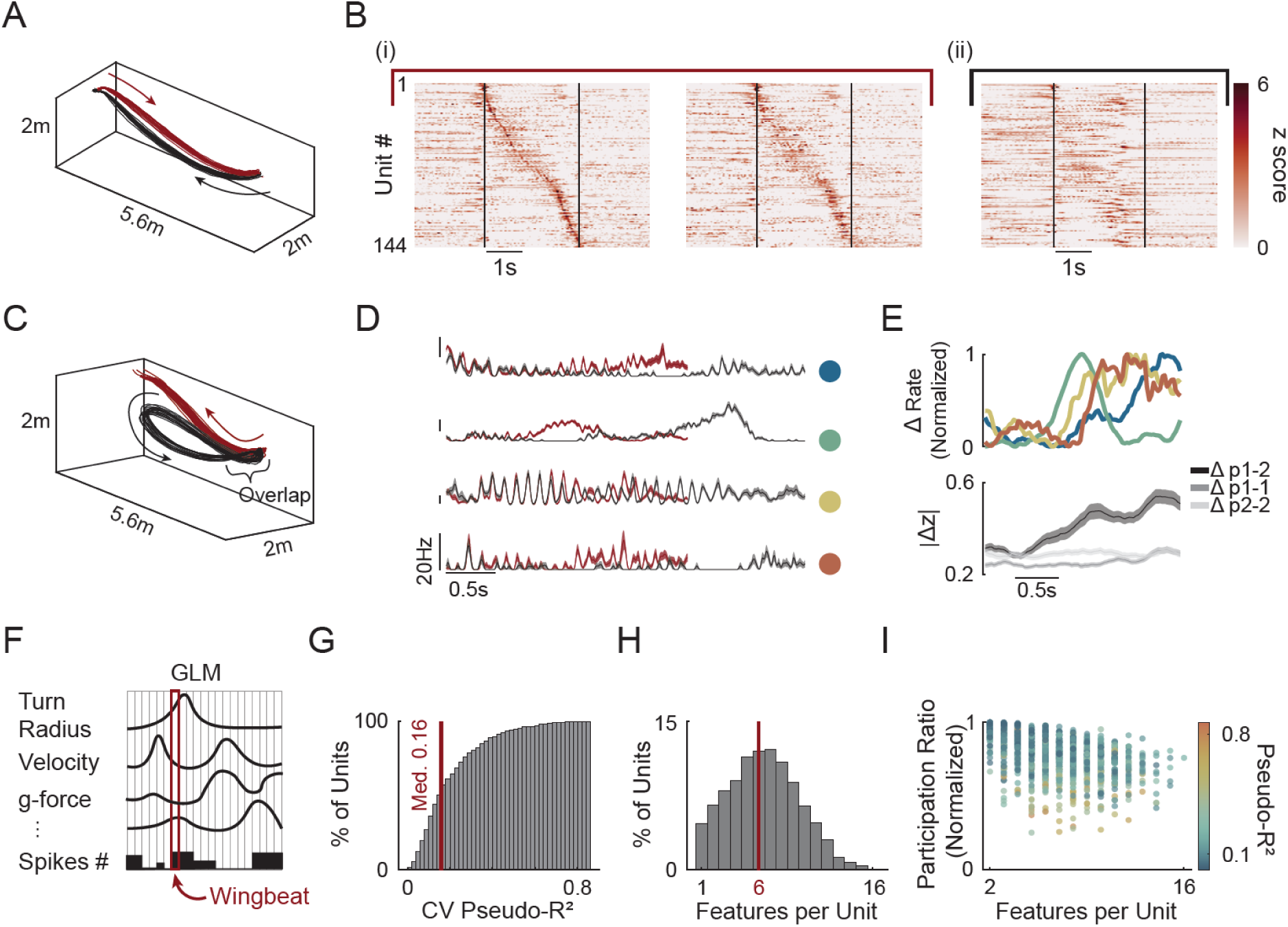
Motor cortical activity tracks kinematic demands across flight paths. **A)** Two example flight paths from the same session with similar duration. Path #1 (feeder to the perch, red) and Path #2 (perch to feeder, black). **B)** Mean z scored, smoothed activity of 144 co-recorded units during the paths shown in (A). (i) Left: odd numbered flights of Path #1 (ordered by time of peak activity). Right: even-numbered flights of the same path, using identical unit order. (ii) All flights of Path #2, ordered as in (i). Vertical black lines denote flight start and end. **C)** Two example paths with overlapping takeoff trajectories that gradually diverge. Path #1 (red) and Path #2 (black). **D)** Mean ± SEM firing-rate traces of four example units across the two paths in (C). Colors correspond to paths in (C). Colored dots denote units referenced in (E). **E)** Mean absolute z-scored difference in activity between the two paths in (C). Top: example units from (D). Bottom: mean ± SD across all units. Dark gray: across-path differences; light gray: within-path differences (odd vs. even flights). Mean calculated using sliding window size of 250 ms in 5 ms steps. **F)** Schematic of data used for training unit specific Poisson-GLMs. Spike number in each wingbeat was fitted against the mean value of each flight kinematic feature during that wingbeat cycle (n = 17 features). Red box marks one example wingbeat. **G)** Distribution of 5-fold cross validated pseudo-R2 values across all units and paths (n = 1,331 units, N = 5 bats; median = 0.16). **H)** Distribution of the number of features selected for each units’ model. Number of features was selected using 5-fold cross validated elastic-net regularization. This selection process was used prior to fitting the units shown (G). **I)** Normalized participation ratio (PR) of GLM weights for each unit, relative to maximal PR given the number of selected features. PR = 1 indicates maximally distributed weights. Color indicates pseudo-R² (Methods).

We next asked whether units activated dynamically in relation to the unfolding kinematic demands rather than reflecting a distinct, but predetermined, sequence. To do so, we compared different flight paths, produced by the same bat, with similar initial kinematics that gradually diverged. For example, one path that looped back to the feeder while the other climbed towards the perch (Figure 3C). If recruitment tracks flight kinematics, then units should show similar activity in the periods when the flights overlap and diverge when they separate. Indeed, individual units exhibited precisely this pattern - activity started nearly identical across paths but rapidly diverged as flight kinematics separated (Figure 3D). Critically, each unit diverged at a different timepoint (Figure 3E, top). Since different features (e.g., speed, acceleration, turn radius) change at different moments as flight paths separate, this might indicate that units have distinct kinematic tuning, diverging when their specific driving features separate. Indeed, at the population level, this produced a gradual accumulation of difference as the flight paths diverged, reaching stable between-path distinction only after the trajectories fully separated (Figure 3E, bottom), consistent with high variable kinematic sensitivity at the population level.

Yet what kinematic features drive selective recruitment? To address this question, we fitted generalized linear models (GLMs; Methods) relating each unit’s activity to flight-level kinematics which are the variables that must be controlled to achieve path reproducibility. Across bats and paths, models captured substantial variance (median CV pseudo-R² = 0.16; IQR = 0.18; n = 1,331 units, N = 5 bats). Notably, the upper quintile showed stronger encoding, with 25% of units exceeding a CV pseudo-R² of 0.27 (333/1,331 units, Figure 3F,G). Importantly, units displayed mixed selectivity with each responding to a distinct combination of kinematic features often with no single kinematic variable dominating across the population (Figure 3H-I). Thus, the selective recruitment emerges from diverse kinematic tuning where each unit is sensitive to a specific multi-dimensional state, potentially involving both sensory input and motor output. This results in the sparse, dynamically recruited population we observed.

### Units show a continuum of wingbeat phase tuning and temporal precision, reaching millisecond-scale accuracy

A key component of an accurate implementation of wingbeat cycle-specific adjustments is the need for high temporal precision, since kinematics adaptations could have differing effects if implemented at different times along the wingbeat cycle^38^. We reasoned that motor cortical activity may reflect the nested timescales of both the slower evolution of each flight pattern and the faster one for each individual wingbeat, potentially integrating the two across the population. Indeed, we found that over a third of the recorded unit/path pairs showed significant bias relative to the wingbeat cycle phase (1621/4489 unit-path pairs; Benjamini–Hochberg FDR, q = 0.05; Figure 4A-C). Across the population we observed a spectrum of temporal precision (Figure 4A-B). At one end we observed units that exhibited only a weak, yet significant, bias towards a specific phase of the wingbeat and on the other extreme, highly selective units that fired only a few spikes per cycle with millisecond-scale jitter for a specific wingbeat group within a path (Figure 4B). As a whole, tuning strength varied across the population, from weak phase bias to tight temporal precision (Figure 4D) and the population of phase-locked units uniformly tiled the wingbeat cycle with a high level of reproducibility (Figure 4E). Furthermore, phase preferences were uniformly distributed across the wingbeat cycle with no clear preference toward the upstroke or the downstroke (Figure 4F).

**Figure 4.**
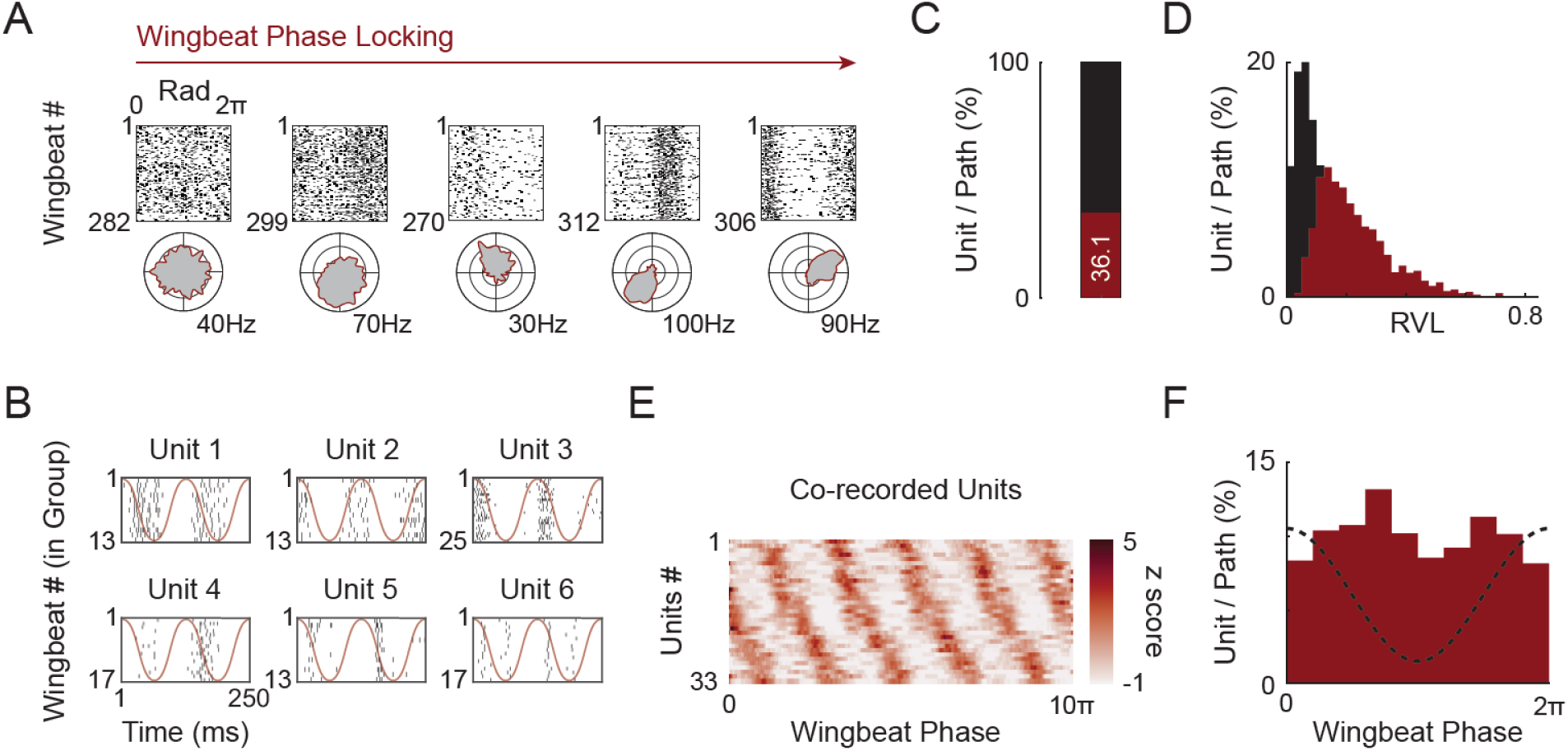
Wingbeat phase bias and temporal precision. **A)** Examples of wingbeat phase–aligned spike rasters for a single path (top) and corresponding polar histograms (bottom). Silent wingbeats are not shown. Units are ordered left to right by increasing phase-locking strength. Outer ring denotes firing rate. **B)** Example raster plots of six units across two consecutive wingbeats (250 ms window). Red line indicates wingbeat phase across the two wingbeats. **C)** Fraction of unit–path pairs exhibiting significant wingbeat phase locking. Each unit was tested separately for each path. Significant locking determined using Benjamini–Hochberg FDR correction (q = 0.05; threshold computed per path). 1,621 of 4,489 (or 36.1%) unit–path pairs were significant. **D)** Distribution of RVL values for all wingbeat-locked (red, n = 1621) and non-locked (black, n = 4489) unit-path pairs. **E)** Mean z scored activity of strongly phase-locked co-recorded units (n = 33 units). Mean activity traces were computed using a sliding window across the whole path (5 wingbeats window size and 4 wingbeat overlap). **F)** Distribution of preferred wingbeat phase (mean angular direction) across all phase-locked unit-path pairs.

Notably, while some units showed varying phase locking properties between paths (Figure S7A, unit B) a subset of phase-locked units remained active throughout the flight, firing at consistent phases of each wingbeat regardless of changing flight demands (Figure S7A unit A). These continuously active units could provide a stable temporal scaffold for aligning more kinematically sensitive units to the correct phase. To test this notion, we evaluated the degree to which the general wingbeat phase can be decoded across different paths. Using least squares regression onto sine and cosine basis functions of the phase, we trained models on the subset of wingbeat-locked, continuously active units in one flight path and tested them on another (Methods). We found that using this subset of units, the general wingbeat phase could be well predicted from one path to another, even when flight kinematics differed markedly (Figure S7B and C, median normed-R^2^ = 0.63; IQR: 0.22; n = 22 path pair comparisons).

These results reveal that a subpopulation of continuously active units exhibits the stable rotational dynamics reminiscent of both single reaches and continuous cycling tasks in primates^10,32^. However, these dynamics do not dominate the population. Instead, the diversity of phase-locking and kinematic sensitivity regimes across units suggests that motor cortex employs flexible temporal strategies to support the millisecond-scale precision required for flight control.

### High dimensionality emerges across wingbeat and is supported by a large shared subspace

Our findings thus far have revealed that co-recorded units exhibited remarkably diverse activity patterns, reflecting the two nested timescales of the wingbeats and flight paths with the evolving timescales of selective recruitment (Figure 5A). This led us to hypothesize that, in contrast to many laboratory tasks where dimensionality does not expand dramatically across different movement conditions^11,31,32,39^, population dynamics in the motor cortex of flying bats will be high-dimensional across different wingbeats (Figure S5).

**Figure 5.**
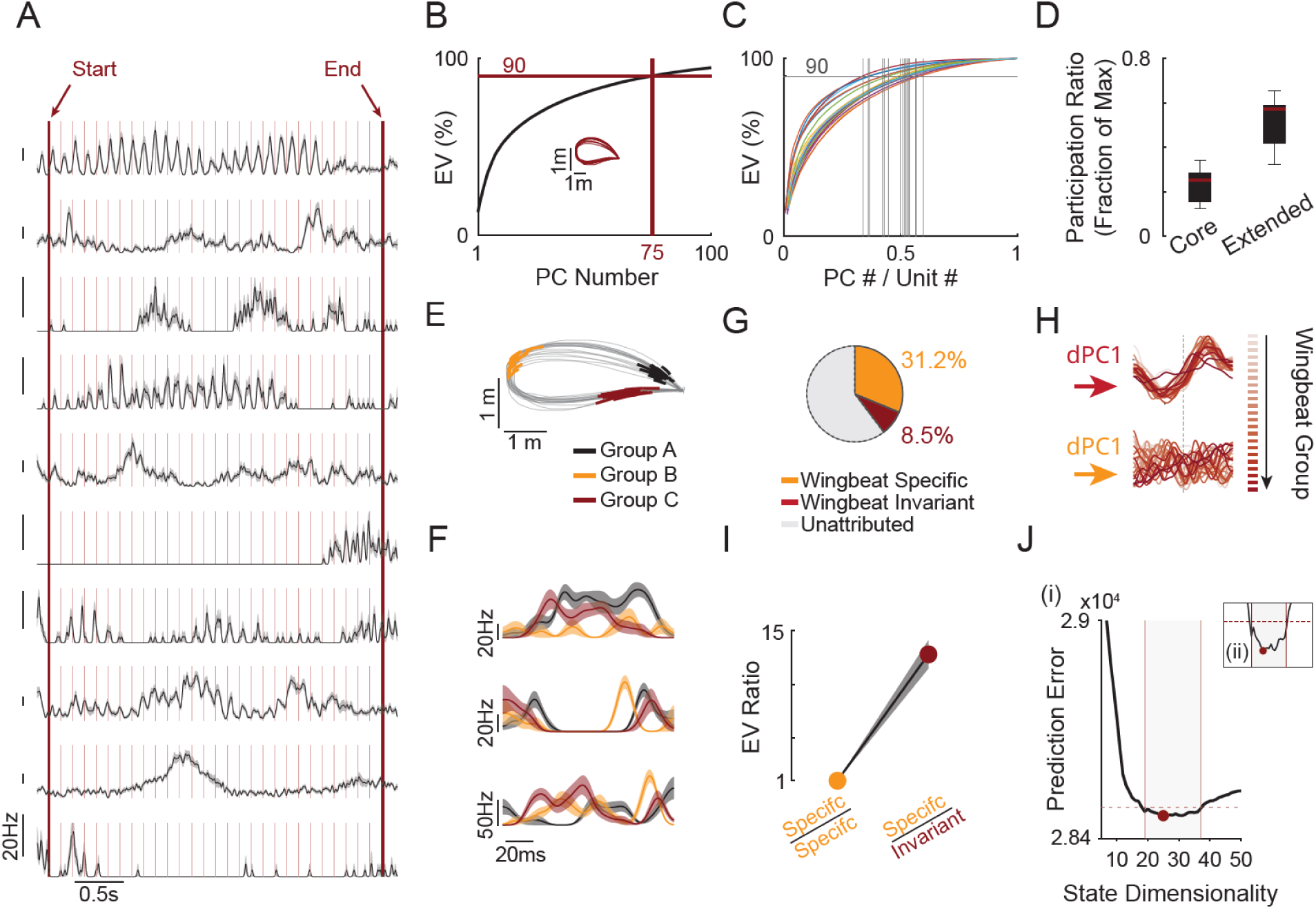
High dimensionality emerges across wingbeats. **A)** Mean ± SEM of nine co-recorded units for an example flight path. Light red lines denote wingbeat boundaries; thick red lines mark the first and last detected wingbeats. **B)** Example of the cumulative explained variance (EV) curve obtained from PCA analysis of one example path (inset). Only the first 100 principal components (PC) are shown. Reaching 90% EV required 75 PCs (N = 203 units). **C)** Cumulative EV curves for all paths normalized to the number of recorded units. Each colored curve represents one path. Gray vertical lines indicate the fraction of components needed to reach 90% EV for each path. Only paths with at least 50 units were used. **D)** Participation ratio across the whole spectrum (“Core”) and for the extended spectrum (components beyond the PC capturing 50% EV). Values normalized to the maximum number of components. Box plots show median, IQR (boxes), and 1.5× IQR (whiskers). **E)** Example flight path with three selected wingbeat groups colored. **F)** Three example units’ (mean ± SEM) activity across the wingbeat groups shown in (E). Colors correspond to wingbeat groups. **G)** Demixed PCA (dPCA) decomposition for the same path as in (E) showing percentage of EV attributed to wingbeat-group–specific components (group and group × time), wingbeat-group–invariant components (time only), and unattributed variance. **H)** First demixed principal component (dPC1) for invariant (top) and group-specific (bottom) marginalizations. Each trace represented one wingbeat group. Colors indicate temporal order along the path. **I)** Ratio of group-specific to group-invariant EV across paths (mean ± SEM: 12.0 ± 6.7, p = 23 paths, N = 5 bats). **J)** (i) Example of dimensionality estimation using cross validated GPFA for one selected path. Shaded area indicates estimated range around minimum (using 1% threshold relative to minimum). (ii) Close-up of the prediction error around minimum for the same path.

To estimate the degree of neural dimensionality, we began with a standard PCA analysis similar to that used in previous studies of motor cortical population dynamics^10,34,40,41^. For each wingbeat group within a flight path, we averaged the smoothed firing rates across all wingbeats to obtain a single mean trace per unit. Group means were then concatenated to form a matrix of units × time bins. Higher noise wingbeat groups and units were removed (Methods). Figure 5B shows the PCA results for one example path consisting of 25 wingbeat group clusters. The cumulative variance curve showed the first few PCs carrying substantial variance (PC1 =13.3%, PC2 = 6.6%). However, the rise of the curve decayed relatively slowly with later PCs. The first 10 PCs only explained about 50% of the variance. The remaining half of variance was distributed across a long tail. Indeed, reaching 90% explained variance (EV) required 75 PCs, far exceeding the dimensionality typically reported for constrained motor tasks such as reaching and cycling tasks, tracing paths in a maze or walking on a treadmill^10,11,32,31,33,42,43^. To compare across sessions and bats, we normalized the number of PCs by the number of recorded units, giving us an interpretable metric that expresses how much of the recorded population’s capacity is utilized. Reaching 90% of the variance required, on average, 48.5 ± 7.7% of the available dimensions (units) across bats and paths (mean ± s.d.; Figure 5C; n = 20 paths with at least 50 flight modulated units), several-fold higher than that typically reported, even when absolute dimensionality is relatively high^44^. The steep initial rise followed by a long, slowly decaying tail in the EV curve raised the following question: does this tail reflect structured neural activity or noise? We computed a participation ratio (PR) that quantifies how uniformly is the variance spread across dimensions. PR can also be computed over any subset of the spectrum, allowing us to separately characterize the “core” subspace (PCs capturing the first 50% of variance) and the “extended” subspace (all remaining PCs). If the extended subspace were isotropic noise, its PR should approach its maximal number of dimensions. An intermediate value would instead suggest that the tail itself is high-dimensional yet structured. Indeed, the max-normalized extended PR showed intermediate values indicating structure beyond the core PCs (51.8 ± 10.9 % mean ± s.d.; Figure 5D). In addition, restricting our PCA analysis to increasingly consistent units preserved the high dimensionality, ruling out the possibility that high dimensionality arises from noisier units (Figure S8A-B). Importantly, noise modeling of PCA dimensionality yielded lower bounds that remained elevated (Figure S8C-F).

We next asked what could be driving this high dimensionality? Our results thus far suggest that units can change their activity between individual wingbeats pointing to a low amount of shared variance across these wingbeats which in turn can explain the high dimensionality we observed (Figure S5). Using demixed PCA (dPCA, ref.^45^) allows the separation of variability between shared (condition invariant) and unique (condition specific) components. dPCA has previously been used to demonstrate that a large portion (49-77%) of variability is shared between different movement conditions, regardless of direction or maze-path taken even with upwards of 27 different conditions^11^, which reduces the additional dimensionality gained when analyzing across movements. To directly quantify this, we applied dPCA to our population activity, grouping marginalizations into condition-specific (wingbeat group) and condition-invariant (time only) components regularized via cross-validation on held-out trials (Methods; ref.^45^). We first illustrate this structure using an example flight path (Figure 5E). Units showed diverse yet consistent activity profile for different wingbeat groups (Figure 5F). We found that condition-invariant components only accounted for 8.5% of the total variance, while the condition-specific components captured a much larger portion (31.2%; Figure 5G). The condition-specific dPC1 varied for each wingbeat group while the condition-invariant dPC1 was strongly consistent across the different wingbeat groups as expected (Figure 5H). On average, the condition-invariant component captured only a fraction of the total variance (3.6% ± 2.6 % mean. ± s.d, n = 23 paths). Indeed, the condition-specific explained variance was 12 times larger on average than the condition-invariant portion (Figure 5I, 12.0 ± 6.7 explained variance ratio, mean ± s.d). Thus, the high dimensionality observed could be driven by the fact that different wingbeats do not share a strong invariant driving factor. Instead, motor cortex produces relatively unique patterns of activity for different wingbeats.

While many units show unique activity patterns, our GLM models suggest units share common driving kinematic features (Figure 3), the wingbeat phase coordinates activity across the population (Figure 4), and the first few PCs capture a large portion of the variance. To specifically quantify the portion of variance that is shared among units, we applied Gaussian Process Factor Analysis (GPFA), which explicitly models private neuronal variance separated from common factors (Factor analysis). Importantly, this method allows for optimization of different time constants for each factor (Gaussian processes), a feature that is helpful in modeling our nested behavioral time scales. To estimate the dimensionality of shared population activity, we used leave-one-neuron-out cross-validated prediction error, selecting the number of GPFA factors that best predicted each unit’s activity from the remaining population^46^. The portion of the dimensionality of the shared subspace varied between paths but was relatively high: up to 28 latent factors were required to optimally predict neuronal activity from the population state (20.2 ± 5.8, mean ± s.d, n = 13 paths; Figure 5J, Figure S8G).

Taken together, these results suggest that the high dimensionality arises from weak repetition of patterns across wingbeats which could be driven by coordination across a large population-wide wingbeat varying set of shared factors.

## Discussion

We found that despite the complexity of controlling multi-joint, articulated, hand-like wings, bats reproduced flight paths with remarkable precision by adjusting wingbeat kinematics within a narrow solution space. Motor cortical activity reflected these features: distinct neuronal populations were selectively recruited on successive wingbeats according to specific kinematic states, while a dedicated subset of units encoded wingbeat phase with up to millisecond precision. Combined, these properties produced a high-dimensional population code, several folds greater than that reported previously in constrained motor tasks^10,11,32,31,33,42,43^. This observation is consistent with theoretical links between neuronal dimensionality and behavioral complexity^17,44,47–49^ and underscores the importance of studying neural circuits under ethologically-valid and complex forms of natural behavior.

What accounts for the high dimensionality of motor cortical activity we observed during flight as compared to other behavioral paradigms? In primate reaching tasks for example, at least half of the variance is similar across movements, such that different actions induce only relatively modest modulations of a shared neural pattern^11,31,34^. Much research has since focused on the low-dimensional neuronal trajectories that emerge^50–52^. This aspect of motor cortex seems paradoxical: if motor cortex is essential for generating dexterous behavior, where small differences in movement can have large functional consequences, why are such differences so weakly represented at the level of population activity? Indeed, several prior studies have emphasized that motor cortical activity should not be confined to low-dimensional manifolds and have suggested that increasing behavioral complexity may reveal richer underlying structure^17,47,48^. More complex behaviors such as grasping suggest higher dimensionality^53^, yet the principles underlying these differences remain unclear. In our dataset, less than 5% of variance was condition-invariant across wingbeats. Thus, rather than reflecting small variations on a largely shared template, each wingbeat is associated with unique population activity pattern, as would be hypothesized from the role of motor cortex in controlling the finer aspects of movement. Consistent with that view, the motor cortex has been shown to be crucial for corrective sub-movements during tongue reaches^35^, and motor cortical activity reflects such corrections during reaching^36,54–56^. Notably, the unique population activity patterns observed on each wingbeat need not exclusively reflect outgoing motor commands; a portion of this condition-specific variance may arise from the integration of sensory feedback, which is itself dynamically and selectively modulated during movement^57,58^.

How can these findings be reconciled with results from more constrained laboratory tasks? Optimal control theory predicts that motor systems primarily correct deviations along task-relevant dimensions (the minimal intervention principle^30^). As such, sets of relatively similar endpoint-focused behaviors may naturally favor the reuse of shared, low-dimensional neural patterns across movements^31^. During high-speed and complex 3D flight, however, accurate performance requires precise control of wingbeat kinematics on every cycle, effectively narrowing the range of permissible solutions. In this regime, minimal intervention still applies, but behavioral demands require distinct adaptations at successive wingbeats, leading motor cortex to generate different sequences over time. This framework is consistent with the suggested role of motor cortex in gait adaptation when traversing an obstacle^16,59,6,43^ yet expands it to a regime in which unique adaptation signals need to be integrated into virtually every locomotion cycle. From this perspective, neural dimensionality may depend strongly on behavioral demands and indeed, expansions in dimensionality emerge even in highly constrained low-dimensional tasks when they require sufficiently diverse and continually precise control strategies^60^.

However, even in those settings the reported neuronal capacity utilized (dimensionality normalized to the number of recorded units) was still significantly lower than that observed in our data, suggesting variability might also be related to within-task complexity. This view of motor cortical computation carries practical implications for neuroprosthetic design and motor rehabilitation through brain–machine interfaces. The minimal number of recording channels needed to capture relevant population variance may be substantially higher for dexterous tasks. Similarly, dimensionality reduction methods that work well for simplified movements may discard information critical for dexterity, which still remains a major challenge in the field^61,62^.

More broadly, our results underscore the value of studying the brain during natural, complex and ethologically relevant behaviors. The hand-like structure of the bats’ wing may reflect convergent evolutionary solutions for precise motor control, or species-specific adaptations. Distinguishing these possibilities will require comparative studies^63,64^. Lastly, our findings suggest that complex actions expose computational demands not apparent in simplified tasks, revealing regimes that would otherwise remain hidden. Capturing the full computational landscape of the mammalian brain will require embracing the richness of behaviors that animals have evolved to perform^65,66^.

## Supporting information

Movie 1 - 3D tracking of bat during flight

## Acknowledgements

We thank the members of the Yartsev laboratory, Mehrdad Jazayeri, Preeya Kahana for discussion and comments on the manuscript. We also thank Madeleine Snyder for her illustration in Figure 1A and the staff of the Office of Laboratory Animal Care for support with animal husbandry and care. This research was supported by the NIH National Institute of Neurological Disorders and Stroke (K99NS14234, to B.S) and The Howard Hughes Medical Institute.

## Author Contributions

B.S. and M.M.Y. designed the research. B.S., K.K.Q. and X.C. performed experiments. B.S. analyzed the data with input from W.A.L. and M.M.Y. The manuscript was written by B.S. and M.M.Y. with input and comments from K.K.Q., X.C. and W.A.L. M.M.Y. supervised the project.

---

**Movie 1: Markerless kinematic tracking of a freely flying bat.** Representative flight behavior is shown from three sequential camera perspectives (front, back, and side views). Colored markers indicate anatomical keypoints on the body and wings, extracted using a custom-trained DeepLabCut model. Video was acquired at 120 Hz and is displayed at 10× slower than real time (0.1× speed).

