## Supplementary material for "Motor Cortical Computations Underlying Natural Dexterous Movement in Freely Flying Bats": Methods and Supplementary Figures

**This file contains:**

1. Materials and Methods
2. Supplementary Figures and Legends
3. Supplementary Table 1

*To whom correspondence should be addressed.

 (B.S) and (M.M.Y)

### Materials and Methods

#### Subjects

Experiments were conducted in five adult male Egyptian fruit bats (Rousettus aegyptiacus; body weight ~150 g). Three bats were implanted with one Neuropixels probe and two bats were implanted with three Neuropixels probes. We collected 16 recording sessions (2–4 sessions per bat), during which neural activity was recorded during free flight. All bats were housed in humidity and temperature-controlled rooms. Following implant surgery, implanted bats were singly housed. Lighting was maintained on a 12 h:12 h reverse light cycle, and all experiments were performed during the animals' active (dark) phase. All procedures were approved by the Institutional Animal Care and Use Committee at the University of California, Berkeley.

#### Surgery

Probe implantation was performed as described previously[28] in two stages, separated by at least of 7 days: (i) implantation of the training cone and (ii) insertion of Neuropixels 1.0 probes.

*Implantation of training cone*

The bat was anaesthetized using an injectable cocktail of ketamine, dexmedetomidine and midazolam. It was then placed on a stereotaxic apparatus (Model 942; Kopf) and provided with a continuous supply of oxygen. Anesthesia was maintained by injections of a cocktail of dexmedetomidine, midazolam, and fentanyl according to anesthesia depth which was continuously monitored by toe-pinch reaction test and by measurements of the bat's breathing rate. Body temperature was measured with a rectal temperature probe and maintained at approximately 35 °C using a regulated heating pad. After proper anesthetic depth was reached, the skull was exposed, and surrounding skin and tissue were retracted, and the exposed skull was cleaned. A ground screw, which consists of a bone screw (19010-00; FST) with stainless-steel wires (203.2 μm coated; A-M Systems) soldered to the screw head served as the ground for each Neuropixels probe (one wire per probe). In two bats the ground screw was inserted in the frontal plate of the skull, and in the three others it was inserted posterior to the recording sites above the hippocampus. Three or four shorter bone screws (M1.59 mm stainless steel) were placed to further strengthen the attachment of the implant to the skull. A circular 1-mm craniotomy was made for each probe insertion point, up to three craniotomies per bat. All craniotomies were made at approximately 11.7 mm anterior to the transverse sinus that runs between the posterior part of the cortex and the cerebellum. For single probes, the craniotomy was located 1.5 mm lateral to the midline on the right hemisphere. For three probes, two craniotomies were done bilaterally at 1.5 mm and a third site was added 2.5 mm from the midline on the right hemisphere. The craniotomy was then sealed with a biocompatible elastomer (Kwik-Sil; World Precision Instruments) to protect the brain surface until probe insertion. A custom 3D-printed cone was positioned and cemented using dental acrylic. The cone was closed with a custom 3D-printed cap. At the end of the surgery, after the bat had fully awoken from the anesthesia, an oral analgesic (Metacam; Boehringer Ingelheim), was administered. Analgesics (three days) and antibiotics (seven days) were given daily until complete recovery. Behavioral training was resumed after the bat was allowed to fully recover from surgery for five days. During training, the weight of the implant was gradually increased over several sessions to allow the bats to slowly adapt to the final implant weight

*Insertion of Neuropixels 1.0 Probes*

Before probe insertion, each Neuropixels 1.0 probe was sharpened at a 20°–30° angle for 15 min using a Microgrinder (EG-45; Narishige), and a single stainless-steel wire (203.2 μm coated; A-M

Systems) was soldered connecting both the ground and reference of the probe. The probe insertion procedure follows the same general surgical practice as described above. The training cone was removed and up to three probes were inserted into pre-existing craniotomies after a durotomy in each. After mounting the Neuropixels 1.0 probe on a stereotaxic arm, the probe shank was coated with fluorescent dye (CM-DiI; Invitrogen C7001) and inserted into the target craniotomy at a rate of around 10–20 μm / s. The probe was then cemented in place using dental acrylic. When the cement had fully cured, the ground wire of the probe was connected to the pre-existing ground screw. After all probes had been inserted, a new 3D-printed cone was positioned and cemented to the skull. Each probe was connected to a connector piece (SpikeGadgets) attached to the top of the cone, which serves both as a protective cap and as the interface between the Neuropixels probes and the wireless SpikeGadgets headstage.

**Electrophysiology Data Acquisition and Spike Sorting**

Recordings began one day after probe insertion and were performed using a SpikeGadgets wireless Neuropixels 1.0 headstage, attached prior to each session to the connector on the implanted cone along with a battery and SD card. A maximum of 384 channels were recorded simultaneously across up to three probes. Recording channels were selected to maximize active channels across . Electrical signals (referenced to the ground screw) in the spike band (600–6,000 Hz) and LFP band (0.5–200 Hz) were amplified 500–1,000× and 125–250×, respectively, and were logged locally to an SD card on the headstage. After each recording session, the headstage was removed and the SD card was retrieved. Recorded data on the SD card were downloaded using a logger dock (SpikeGadgets). Drift correction and spike sorting were done automatically using Kilosort4[67]. All units were further manually curated using Phy[68].

**Histology**

At the end of experiment, bats were euthanized with an overdose of sodium pentobarbital and transcardially perfused with 200 ml phosphate-buffered saline (PBS; 0.025 M, pH 7.4), followed by 200 ml fixative (3.7% formaldehyde in PBS). The implant was then carefully removed, and the brain was dissected and stored in fixative. The fixed brain was subsequently moved to a 30% sucrose solution in PBS for cryoprotection, and 40 to 60 μm coronal sections were cut using a microtome (HM450; Thermo Fisher Scientific) with a freezing stage. Slices were stained for DAPI (Thermo Fisher Scientific) and cover-slipped with aqueous mounting medium (ProLong Gold Antifade Mountant, Thermo Fisher Scientific). Fluorescent images of the slides were acquired using a slide scanner (Zeiss Axio Scan 7 Slide Scanner) and used to visualize Neuropixels probe tracks from CM-DiI fluorescence (Invitrogen C7001).

**Flight Room Setup**

All experiments took place in an indoor flight room ( 5.6m × 5.2m × 2.5m) which was acoustically, electrically and radio-frequency shielded. To direct the bat to fly in the area captured by pose tracking cameras (3D pose tracking), several barriers were placed in the middle of the room dividing the room roughly in half. The flight room volume was otherwise empty (except for one session in which obstacles were placed to increase path diversity). For all experiments, periodic clock pulses generated by a Master-9 device (A.M.P.I.) were used to create a timing signature that served as a common frame of reference for all the recording systems (positional tracking using motion capture cameras, 3D pose estimation cameras and neural recording). Daily training and recording sessions consisted of a ~60-90 min period in which the bats engaged in spontaneous flights in the flight room. Bats were mildly food-restricted (>85% of their baseline weight) to motivate flight to a feeder platform which dispensed a small reward for each landing as described previously[29].

**Positional Tracking, Flight Segmentation and Clustering**

The bat's three-dimensional spatial position was tracked at millimeter resolution using 16 motion-capture cameras (ref. [28,29]; Raptor-12HS, Motion Analysis). Each camera detected three infrared reflective markers mounted on the neural recording headstage. The centroid of the marker set was computed using commercially available software (Cortex-64; Motion Analysis).

*Flight Segmentation*

Flight segmentation was done as described previously[29]. Raw marker positions were first preprocessed by taking the median across all markers. Missing values were filled using modified Akima interpolation. The continuous marker position vector was smoothed with a moving median filter (window = 30 frames, 0.25 s) and a Gaussian filter (kernel length = 3 frames). Instantaneous velocity was computed as the Euclidean norm of the frame-to-frame displacement divided by the inter-frame interval. Samples with speed below 1 m/s were classified as non-flying. Flight episodes were identified as sustained periods (≥1 s) of above-threshold velocity.

*Flight Clustering*

Clustering flights into "path" clusters was done as described previously[29]. Each flight was resampled to 20 equidistant points using cubic spline interpolation, yielding a 60-dimensional feature vector (20 points × 3 spatial coordinates) per flight. Agglomerative hierarchical clustering was performed on these feature vectors using Euclidean distance and single linkage. The linkage distance cutoff was manually optimized for each session to ensure spatially consistent flights within each path. Clusters containing fewer than 3 flights were merged into a category of un-clustered flights. Finally, a further step of manual curation was done to ensure the resulting clusters consisted of highly similar flight paths.

**Wingbeat Segmentation**

For detection of the wingbeats, we used accelerometer data acquired at 30 kHz by the neural recording headstage. For each flight time as detected from the position data, the raw accelerometer signal was extracted (± 1.5 s around flight) at the z axis and was bandpass filtered (2–20 Hz, 2nd-order Butterworth, zero-phase) to remove low-frequency drift and high-frequency noise outside the wingbeat frequency range. The signal was then z-scored and a low-frequency trend (2 Hz low-pass, 2nd-order Butterworth) was subtracted to remove residual slow variation. A Gaussian smoothing kernel (window = 33 ms, $\sigma$ = 6.7 ms) was applied to reduce noise. Individual wingbeat peaks were identified using a peak-detection algorithm (MATLAB findpeaks) requiring a minimum peak height of 0.5 z-score units, and a minimum peak prominence of 0.5 z-score units. Each detected peak was assigned a timestamp by mapping the accelerometer sample index to the global recording clock. For each flight, wingbeat durations were computed as the difference between consecutive peak timestamps. Detected wingbeats were further screened to exclude spurious detections. For initial characterization of the wingbeat period distribution (Figure S4), wingbeats were flagged if their preceding interval fell outside [0.5, 2.0] × the flight's median inter-wingbeat interval, using broad limits to capture the full range of observed periods while excluding gross artifacts. For all subsequent analyses, stricter limits of [0.9, 1.1] × the flight's median were applied. Additionally, the 3D Euclidean distance traveled between consecutive wingbeat positions was computed and outliers were identified using a Hampel filter (half-window = 3 wingbeats, threshold = 3 MADs). Wingbeats flagged by either criterion were excluded from all subsequent analyses. Short flights containing fewer than 12 wingbeats after exclusion were discarded entirely.

#### 3D Pose Tracking

Nine high resolution high frame rate cameras (Balser a2A1920-160umBAS) were positioned along the flight path to capture detailed wing kinematics (Figure S1A). Video was acquired at 100 Hz, with each frame triggered by a TTL pulse generated by a Master-9 device to ensure synchronization with the neural and motion-capture recording systems.

##### *Model training*

We used DeepLabCut[27] (version 3.0.0rc6) to obtain 2D key point detections from each camera from a set of 20 key points (see Figure 1H. in total 23 key points were annotated but the head and left/right eyes were not used in any later analysis). We then used Aniposelib[69] to obtain 3D key point estimates via RANSAC triangulation. We labeled 3850 frames with a 23 key point skeleton. 95% (3657 frames) were used for training and 5% (193 frames) were held out for validation. We trained a DLC model with a DEKR-W32 backbone for 625 epochs with a batch size of 4 using 4x L40 (48 GB) GPUs on the SAVIO computing cluster with input image sizes ranging from 1600x1200 to 1920x1200. The best checkpoint was selected based on the mAP on the validation set (epoch 470, mAP = 91.58%, mAR = 92.87%, MAE = 4.89 pixels) and was used for all subsequent inference of 2D key point positions across the dataset. Camera intrinsic and extrinsic parameters were calibrated with a ChArUco board using Multical (RMS 0.54 +- 0.07 pixels). Finally, we used RANSAC triangulation from the Aniposelib library to triangulate the 3D positions of each key point (minimum DLC prediction confidence of 0.1) using subsets of camera views with minimum reprojection error (minimum of 3 cameras).

*Coordinate Transformation from Global to Body-Centered Reference Frame*

Following 3D pose reconstruction, key point coordinates were transformed from the global (room) reference frame to a body-centered reference frame. Reprojection errors were converted from pixels to metric distances using camera calibration parameters. Key points with reprojection errors exceeding 5 cm were excluded. Remaining key points were smoothed with a median filter, and missing values were interpolated using cubic spline interpolation. For all key points except the body frame (left/right shoulders and left/right hips), interpolation was restricted to gaps of ≤5 frames.

The body-centered coordinate system was defined using the four torso key points. The origin was set as the center of mass (centroid of the two shoulder and two hip positions at each time step). To construct the body axes, the lateral (Y) axis was computed as the average of the left-minus-right shoulder and left-minus-right hip unit vectors. The longitudinal (X) axis was defined as the shoulder midpoint minus the hip midpoint, projected onto the plane perpendicular to Y and normalized. The dorsoventral (Z) axis was obtained as the cross product of X and Y. Frame-to-frame consistency of each axis was enforced by flipping any axis whose sign reversed relative to the previous frame. The resulting rotation matrices were converted to unit quaternions, and temporal smoothing was performed using a bidirectional SLERP-based filter. Each key point was transformed into the body-centered frame by applying the inverse of the 4×4 rigid-body transformation matrix (composed of the smoothed rotation and the translation to the body origin) at each frame

*Calculation of Key Point Displacement, Envelopes and Asymmetries*

For each wing key point (n = 16, excluding the four torso landmarks used to define the body frame), the body-frame displacement along each axis (X, Y, Z) was extracted as a time series for every flight. Missing values were filled using spline interpolation over gaps of ≤8 frames. Flights were temporally aligned using a template-matching procedure. For a given axis, each flight in turn served as a reference template, and all other flights were shifted in time to maximize the Pearson correlation with that template across a range of integer sample lags (with maximum temporal shift of two wingbeat durations). The reference flight yielding the highest mean pairwise correlation across all flights was selected, and the corresponding lags were applied uniformly to all three displacement axes (X, Y, Z) to preserve their temporal relationship.

Upper and lower envelopes of each displacement trace were computed to capture the flight varying key point amplitude. To robustly detect peaks and troughs across key points with different displacement scales, each signal was first range-normalized, mean-centered, and detrended by subtracting a 2 Hz low-pass filtered version (2nd-order Butterworth). Peaks (and troughs, by sign inversion) were identified on this processed signal using a minimum inter-peak distance of 8 samples (~80 ms). The corresponding peak and trough time stamps were used to extract the displacement signal from the original (non-detrended) signal. Upper and lower envelopes were obtained by linearly interpolating through these values across time. Envelopes with insufficient variation (median within-flight standard deviation $< 1.5$ cm) were excluded from subsequent analyses, to avoid estimates dominated by noise.

Bilateral asymmetries were computed for three homologous key point pairs (left/right elbows, wrists, and wingtips (3rd digit, 3rd joint)) by subtracting the left envelope from the right envelope after Gaussian smoothing ($\sigma = 12$ samples). Asymmetry features were only retained when both the left and right envelopes of a pair independently exceeded the 1.5 cm variability threshold. To obtain the distribution of displacement or envelope and asymmetry correlations we computed the correlation for each feature to the same feature across all other flights of the same path. This was then pooled across paths and features.

**Position-derived Kinematic Features**

Flight kinematics were derived from the three-dimensional position of a head-mounted marker tracked by the motion-capture system. For each flight, kinematic measures were computed only over the interval between the first and last detected wingbeats. All derivative quantities were computed using central differences and subsequently zero-phase filtered with a 2nd-order Butterworth low-pass filter (cutoff: 3 Hz), applied via forward-backward filtering to eliminate phase distortion.

The following kinematic features were extracted for each flight. Flight speed was computed as the Euclidean norm of the 3D velocity vector, with velocity obtained as the numerical derivative of position. The three velocity components ($v_x$, $v_y$, $v_z$) were retained separately. Acceleration was computed as the numerical derivative of the filtered velocity vector and similarly low-pass filtered;

the three acceleration components ($a_x$, $a_y$, $a_z$) were retained. G-force was defined as the magnitude of the non-gravitational acceleration, computed by subtracting the gravitational vector [0, 0, −9.81] $m/s^2$ from the acceleration vector and normalizing by g. Flight path angle was defined as the angle between the velocity vector and the horizontal plane: $\gamma = \arcsin(v_z / \text{speed})$. Angular velocity was derived from path curvature, computed as $\kappa = \|\mathbf{v} \times \mathbf{a}\| / \|\mathbf{v}\|^3$, with angular velocity given by $\omega = \kappa \cdot$ speed; a speed-dependent weight (min(speed / 1.5 m/s, 1)) was applied to suppress unreliable curvature estimates at low speeds. Turn radius was computed directly from curvature as $R = 1/\kappa$, capped between 0.1 and 50 m to exclude physically unreliable values. Jerk was computed as the numerical derivative of the filtered acceleration vector and low-pass filtered; both the three components ($j_x$, $j_y$, $j_z$) and the Euclidean magnitude were retained. Mechanical power was estimated as the rate of change of total mechanical energy, $P = d(\frac{1}{2}mv^2 + mgz)/dt$, where m is bat mass (estimated as 145 g), v is flight speed, and z is vertical position, reflecting the net power required to modulate the bat's kinetic and potential energy over the course of each flight.

To obtain the distribution of kinematic feature correlations, flights were temporally aligned using the same template-matching procedure described above (*Calculation of Key Point Displacement, Envelopes and Asymmetries*). We then computed the correlation for each feature to the same feature across all other flights of the same path. This was then pooled across paths and features.

**Wingbeat Adaptation Cosine Similarity**

We computed the cosine similarity between wingbeat-averaged kinematic vectors as a function of wingbeat lag. For each flight, we constructed a feature matrix (timepoints × features) comprising the upper and lower kinematic envelopes of each tracked key point (smoothed with a 6-sample moving average) and bilateral asymmetry envelopes for selected key point pairs (smoothed with a 12-sample Gaussian kernel), as described above. The feature matrix was segmented into non-overlapping windows of 12 samples (one wingbeat cycle), and features with envelope durations shorter than 70% of the median duration across features were further excluded. Each feature was z-scored within flight, and each wingbeat window was summarized by its temporal mean, yielding a single state vector per wingbeat cycle. Pairwise cosine similarity was computed between all wingbeat state vectors within a flight, and values were grouped by wingbeat lag. Flights with fewer than 5 valid wingbeats were excluded.

**Wingbeat Group Clustering**

We clustered individual wingbeats within a path based on their kinematic features using a two-stage dimensionality reduction and clustering approach. For each wingbeat, kinematic features were averaged across the wingbeat duration to yield a single feature vector per wingbeat (a "time in flight" feature was added to the set of kinematic features to improve the temporal aspect of the clustering). All features were normalized to the range [0,1] prior to further analysis. Wingbeat feature vectors from all flights of a given path were then concatenated into a single matrix and embedded into a 10-dimensional space using UMAP Projection with Euclidean distance as the metric. UMAP hyperparameters (minimum distance: 0.1–0.85 in steps of 0.1; number of neighbors: 20–110 in steps 15) were optimized via grid search to yield wingbeat groups that minimized the number of repeated group labels within flight. This yielded groups that closely matched the number of cycles in a path and preserved their temporal order. The advantage of this approach is that it captures even highly selective units active for just one cycle and allows averaging across flights by concatenating activity of individual wingbeat groups. Thus, for each UMAP parameter combination we computed the score $S$:

$$S = \mu \left( \frac{|u_f|}{N_f} \right) - \frac{1}{2} \sigma \left( \frac{|u_f|}{N_f} \right)$$

where $| \mathcal{U}_f |$ is the number of unique cluster labels in flight $f$, $N_f$ is the total number of wingbeats in flight $f$, and $\mu$ and $\sigma$ denote the mean and standard deviation across flights. The standard deviation term penalizes solutions with high variance in cluster diversity across flights. The optimal hyperparameters maximize $S$.

**Flight Active and Flight Modulation of Single Units**

Units were classified as flight-active if they fired ≥2 spikes in at least 70% of flights for at least one path. To identify flight-modulated units, we tested whether each active unit showed consistent firing patterns across flights. Firing rates were smoothed with a 3-bin moving average (15 ms). For each flight, we computed the Pearson correlation between that flight's firing pattern and the mean pattern of all other flights (leave-one-out). The mean correlation across flights quantified pattern consistency. Statistical significance was assessed by comparing observed correlations to a null distribution generated by circularly shifting each flight's firing rate by a random amount (between 2 wingbeat cycles and the flight duration minus 2 cycles; 300 iterations). This preserved temporal autocorrelation while disrupting consistent temporal structure. P-values were computed from z-scores and corrected for multiple comparisons using Benjamini-Hochberg FDR (q = 0.01). Units passing FDR correction in at least one path were classified as flight-modulated.

**Silent Wingbeats and Wingbeat Groups**

For each unit, we counted the number of valid wingbeats containing at least one spike across all flights in a session and divided by the total number of valid wingbeats, yielding an active wingbeat ratio. The silent wingbeat ratio was defined as 1 minus this value. Units with a session-wide mean firing rate below 0.1 Hz or an active wingbeat ratio below 0.05 were excluded from further analysis to avoid inflating sparsity estimates.

We separately computed silent ratio for UMAP clustered wingbeat groups. For each path with at least 8 flights, we determined whether a unit was active in each wingbeat group by computing the fraction of wingbeats in that group containing at least one spike. A unit was considered active in a wingbeat group if it fired on at least 65% of the wingbeats in the group. The silent wingbeat group ratio was defined as the fraction of wingbeat groups in which the unit did not meet this criterion.

**SVM Wingbeat Group Classifier**

We trained a multiclass support vector machine (SVM) to decode wingbeat number from neural population vectors. For each wingbeat, we constructed a binary vector indicating whether each unit fired at least one spike. We used a linear kernel SVM with one-vs-one multiclass coding (ECOC) and uniform class priors. Decoder performance was evaluated using 5-fold stratified cross-validation. We computed balanced accuracy (mean per-class recall) as well as band accuracy, which measures the proportion of predictions falling within ± k wingbeats of the true position (k = 1, 2, 3). Statistical significance was assessed via permutation testing (100 iterations), shuffling wingbeat labels to generate a null distribution. Analysis was restricted to recording sessions with ≥100 units and ≥10 flights per path.

**Neural Population Cosine similarity Across Wingbeats**

We computed the cosine similarity between population activity vectors at varying wingbeat lags. For each neuron, we computed the mean firing rate across all wingbeats and subtracted it from the spike count vector, yielding a mean-centered activity matrix. Neurons with an active ratio (fraction of wingbeats with at least one spike) exceeding 0.9 were excluded to remove tonically active units. For each flight, we computed the cosine similarity between all pairs of population vectors separated by a given wingbeat lag (1–8 wingbeats). Similarity values were averaged across all valid pairs within each flight, producing one decay curve per flight.

**GLMs**

We fit Poisson generalized linear models (GLMs) predicting single-unit spike counts per wingbeat from kinematic features. For each unit, we computed spike counts per wingbeat across all wingbeat groups and paths within a recording session. We extracted the following kinematic features per wingbeat: velocity components ($v_x$, $v_y$, $v_z$), speed, acceleration components ($a_x$, $a_y$, $a_z$), g-force, angular velocity, flight path angle, turn radius, jerk components ($j_x$, $j_y$, $j_z$), total jerk, mechanical power, and time within flight. Prior to model fitting, we removed multicollinear features in two steps: first, we iteratively removed features from pairs with absolute Pearson correlation exceeding 0.9, preferentially dropping the feature with higher mean absolute correlation to the remaining set; second, we iteratively removed the feature with the highest variance inflation factor (VIF) until all

VIFs fell below 5, with a minimum of 5 features retained. This pruning was performed independently for each session. We then used elastic net regularization (α = 0.5) with a log link function to identify kinematic features predictive of firing rate. The regularization parameter λ was selected via 5-fold cross-validation, using the largest λ within one standard error of minimum deviance to favor parsimonious models. Features with non-zero coefficients were retained. We assessed model performance using 5-fold cross-validation only on the subset of selected features, computing pseudo-$R^2$ (1 − deviance_model / deviance_null) on held-out data, where the null model prediction was the mean spike count of the training fold.

To characterize how GLM coefficients were distributed across selected features for each unit, we computed a normalized participation ratio. The participation ratio measures how uniformly a set of values are spread. When applied to coefficient vectors, it captures whether a unit's firing rate is dominated by a single kinematic feature or shaped by many features with comparable influence. Since the participation ratio's raw value scales with the number of elements and each unit had a different number of selected features after elastic net, we normalized by $k$ (number of selected features) to obtain a metric comparable across units. Thus, for a unit with $k$ selected features and coefficient vector **w**, we defined:

$$PR_{norm} = \frac{1}{k} \cdot \frac{\left(\sum_{i=1}^{k} |w_i|\right)^2}{\sum_{i=1}^{k} w_i^2}$$

This metric ranges from $1/k$ (all weight concentrated on a single feature) to 1 (weight spread equally across all features). Units with negative cross-validated pseudo-$R^2$ or no selected features were excluded from this analysis.

**Wingbeat Phase Locking and Decoding**

We computed circular statistics for each unit/path pair separately. Each spike was assigned a wingbeat phase (0–2π) by interpolating its time within the corresponding wingbeat cycle. Only unit/path pairs with at least 25 spikes were included in the analysis. For each valid unit/path pair, we computed the mean resultant vector length (RVL) and mean phase direction from the distribution of spike phases for a specific path, using the Circular Statistics Toolbox in MATLAB[70]. Statistical significance of phase locking was assessed using the Rayleigh test for non-uniformity. To correct for multiple comparisons across all unit/path pairs, we applied the Benjamini-Hochberg false discovery rate (FDR) procedure at $q = 0.05$.

*Cross-path Wingbeat Phase Decoding*

We trained linear decoders on mean phase matrix of population activity from one path and evaluated on activity in a different path in the same session. Phase matrix resolution was set to 25 bins per cycle (~5 ms / bin). We only selected units showing phase tuning (Rayleigh vector length ≥ 0.1) on either path and consistent activity (active on ≥ 50% of wingbeats) on both paths. Activity was normalized per unit by standard deviation. Decoder weights were fit using ordinary least squares regression for the sine and cosine of the phase separately. Decoder performance was quantified as $R^2$ between predicted and true wingbeat phase components. To account for noise ceiling, cross-path $R^2$ was normalized by within-path $R^2$ (train and test on same path).

**Dimensionality Analysis**

*Preparation of Mean Activity Matrix.*

For each path, we constructed a firing rate matrix for dimensionality analysis. Spike times were assigned to phase bins based on their position within the wingbeat cycle (0 to 2π), using 125 equally spaced bins spanning the full cycle. The binned spike counts were then convolved with a Gaussian kernel (σ = 5 ms; resulting FWHM of 11.8 ms, equivalent to ~10% of the wingbeat cycle). This kernel size preserves the fine-scale temporal precision evident in single unit rasters while providing sufficient smoothing for rate estimation across the wingbeat cycle (see Figure 4B).

For each wingbeat group, we averaged the smoothed firing rates across all wingbeats to obtain a single mean trace per unit. Importantly, wingbeat groups with low rates (in which fewer than 70% of wingbeats contained at least one spike) were zeroed to avoid noise-dominated averages. Group means were then concatenated to form a matrix of units × phase bins. Units for which fewer than a threshold of wingbeat groups passed a consistency test were excluded from further analysis (see Spike Count Consistency Within Wingbeat Groups for specifics). The firing rate matrix was soft-normalized using a common procedure[32] . Specifically, we divided each unit's activity by its firing rate range plus 5 Hz, balancing the contribution of high- and low-dynamic-range units while preventing amplification of weak responses. This matrix was used for PCA and dPCA analysis.

*Spike Count Consistency Within Wingbeat Groups*

We evaluated the consistency of spike counts within each wingbeat group. For each unit and each wingbeat group, we counted the number of spikes per wingbeat across all flights. Under the assumption of a stable underlying firing rate, these spike counts should follow a Poisson distribution with a Fano factor (FF) close to 1. We tested this using a chi-squared test against the null hypothesis that spike counts are compatible with Poisson variability ($p > 0.05$). For each unit, we computed the fraction of its wingbeat groups that passed this consistency test. This fraction (FF_ratio) quantifies how reliably a unit's firing follows the expected statistical structure across different wingbeat group clusters. A default value of FF_ratio ≥ 0.5 was used as threshold across analysis unless otherwise specified.

*Dimensionality Estimation using PCA*

Principal component analysis (PCA) was applied to the wingbeat group concatenated firing rate matrix mentioned above. We quantified dimensionality using three metrics. First, EV90: the number of principal components required to explain 90% of total variance. Second, participation ratio: $PR = (\Sigma\lambda)^2 / \Sigma\lambda^2$, where $\lambda$ are the eigenvalues. Third, extended participation ratio: eigenvalues beyond the 50% cumulative variance threshold were isolated, renormalized to sum to 1, and participation ratio was computed on this subset. Isometric noise would produce a normalized extended participation ratio reaching close to the maximal number of eigen values available. A structured high dimensional tail on the other hand would instead reach a substantial but lower number as was the case (Figure 5D). To enable comparison across populations of different sizes, all metrics were also expressed relative to the number of units. To ensure this normalization does not inflate dimensionality estimations, this analysis was restricted to paths with ≥50 flight-modulated units.

*PCA Dimensionality as a Function of unit Consistency*

We performed a control analysis examining how dimensionality metrics change when progressively restricting the population to more consistent units. This allowed us to check if the most consistent low-noise units still retain high diversity of activity profiles, or if low-noise units display strong shared variance. We varied the consistency threshold (FF_ratio) from 0.3 to 0.85, including only units whose consistency ratio exceeded FF_ratio. For each threshold, we computed the relative EV90 (number of principal components explaining 90% of variance, divided by the number of units).

*Noise Estimation of PCA Dimensionality with Flight Number*

To obtain robust dimensionality estimates from PCA analysis, we used a subsampling approach to extrapolate how EV90 stabilizes with the number of flights in each path. For each value of $k$ from 1 to $F$ (where $F$ is the total number of flights), we randomly selected $k$ flights, constructed the neural activity matrix using only wingbeats from those flights, and computed EV90. We repeated this procedure 30 times per $k$ (or used all combinations when fewer than 30 existed) and took the median EV90 across repetitions. EV90 decreases with increasing $k$ as trial-averaging reduces noise. To estimate the asymptotic dimensionality, we fit an exponential decay model:

$$EV90(k) = d + A * e^{-\frac{k-1}{\tau}}$$

where $d$ is the true dimensionality, $A$ is the noise amplitude, and $\tau$ is the decay timescale in flights. We fit this model using weighted least squares, with weights proportional to the inverse variance of the median at each $k$. The asymptotic dimensionality d was estimated as the model's asymptote, and we defined convergence as the number of flights $k_{90}$ at which the noise term decays to 10% of d.

*Demixed PCA*

We applied demixed principal component analysis (dPCA) using the dPCA toolbox (https://github.com/machenslab/dPCA). Firing rates were computed for every wingbeat group separately using the same parameters as above (125 phase bins / wingbeat group, σ = 5 ms, soft-normed, and sub selecting units by 0.5 FFth). This yielded a 4D tensor of dimensions N × G × T × K, where N is the number of units, G is the number of wingbeat groups, T = 125 phase bins, and K is the number of trials (wingbeats) per group. Variance decomposition: We defined two marginalizations: condition-specific variance (wingbeat group main effect + wingbeat group × time interaction) and condition-invariant variance (time main effect only). The regularization parameter λ was optimized via cross-validation (50 repetitions), and noise covariance was estimated from single-trial data. We extracted up to 50 dPCA components (or N−1 if fewer units were available). Sessions with fewer than 30 units were excluded from the analysis.

*Gaussian Process Factor Analysis (GPFA)*

We applied Gaussian-Process Factor Analysis (GPFA) using the publicly available MATLAB toolbox[46] (version 3.00; https://users.ece.cmu.edu/~byronyu/software.shtml). We first constructed a spike count matrix with 1 ms bin size for each flight. Each path was tested separately. We set binWidth to 20 ms and xDims to 50. To determine the optimal latent dimensionality, we performed leave-one-neuron-out 5-fold cross-validated prediction error, selecting the number of GPFA factors that best predicted each unit's activity from the remaining population as described previously[46].

#### Supporting Supplementary Figure

Figure S1. 3D pose estimation model

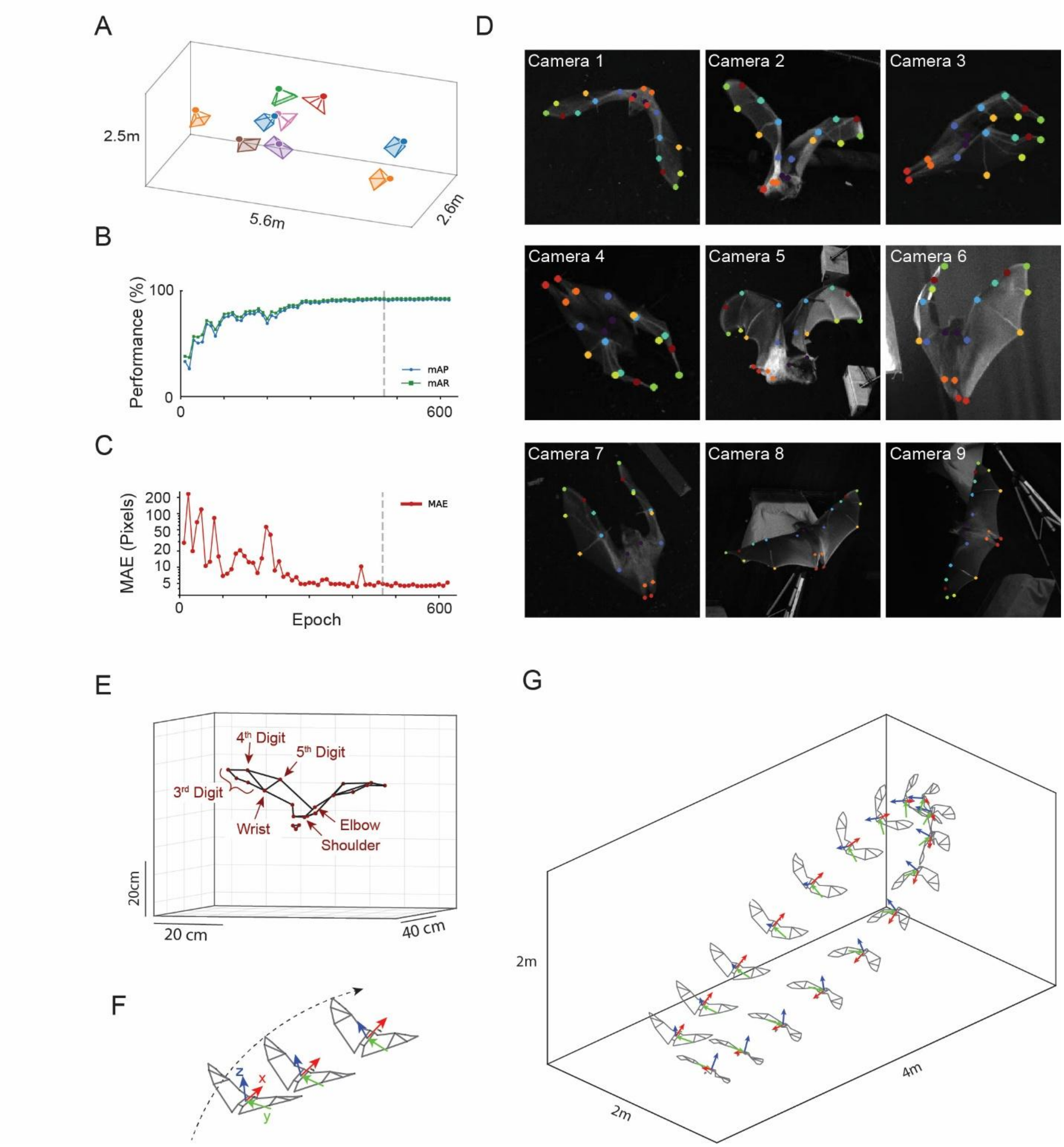


**Figure S1. 3D pose estimation model. A)** Calibrated camera configuration for a typical recording session. Colored pyramids indicate the position and orientation of the nine cameras in the flight room. **B-C)** Model performance during training on a held-out validation set. (B) Mean average precision (mAP, blue) and recall (mAR, green). (C) Mean average error (MAE) evaluated every 10 epochs. Dashed line indicates the selected checkpoint (highest validation mAP) used for pose estimation of the full dataset. **D)** Single frame, keypoint detections from the nine cameras. Each frame captured during different epochs of flight to display array of typical poses. Each dot represents one key point detection from the DLC model. Colors denote matching key points across the left and right of the bat's body. **E)** Reconstructed 3D pose from multi-camera detections (Methods). Several example key points are indicated. **F)** Body-centered reference frame computed from the four torso keypoints (bilateral shoulders and hips). Longitudinal (X, red), lateral (Y, green), and dorsoventral (Z, blue) axes are shown for three frames spaced 12 frames apart, illustrating how the body coordinate system rotates with the bat during flight. **G)** Body reference frame (every 12th frame) along a flight trajectory in room coordinates. Note the rotation of the body axes during the turn relative to the fixed room frame.

Figure S2. Targeting wing motor cortex

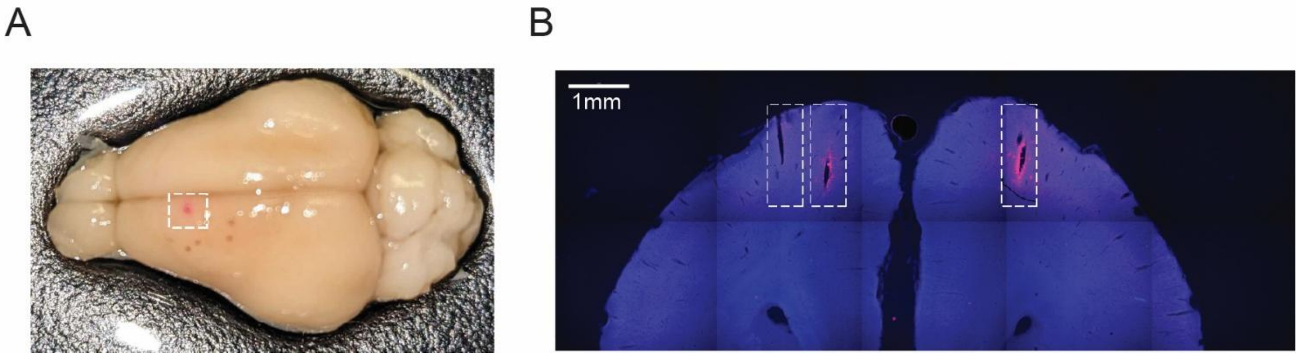


**Figure S2. Wing motor cortex in Egyptian fruit bat. A)** Site of intracortical microstimulation eliciting wing response is indicated using fluorescent dye (white rectangle; following previously published work (Halley et al., 2022)). **B)** Coronal section showing Neuropixels probe tracks (CM-DiI, red) and DAPI staining (blue). Dashed box indicates probe trajectory.

Figure S3. Flight kinematics during free flight

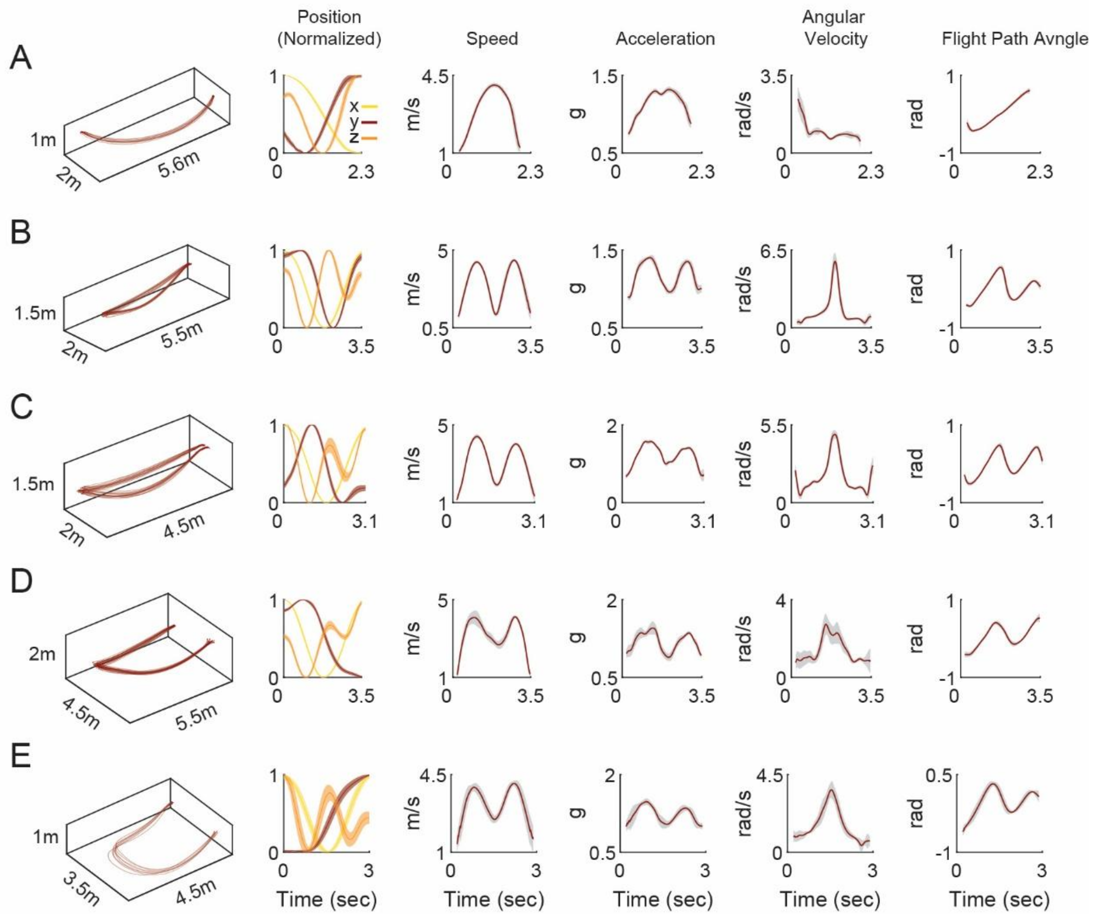


**Figure S3. Flight kinematics during free flight. A-E)** Five example flight paths. Left panel: 3D positions. Right panels: range normalized positions (0-1) and a set of four kinematic variables: speed, g-force, angular velocity and flight path angle. Curves show mean ± SD across flights of the same path.

Figure S4. Wingbeat period variability and adaptation

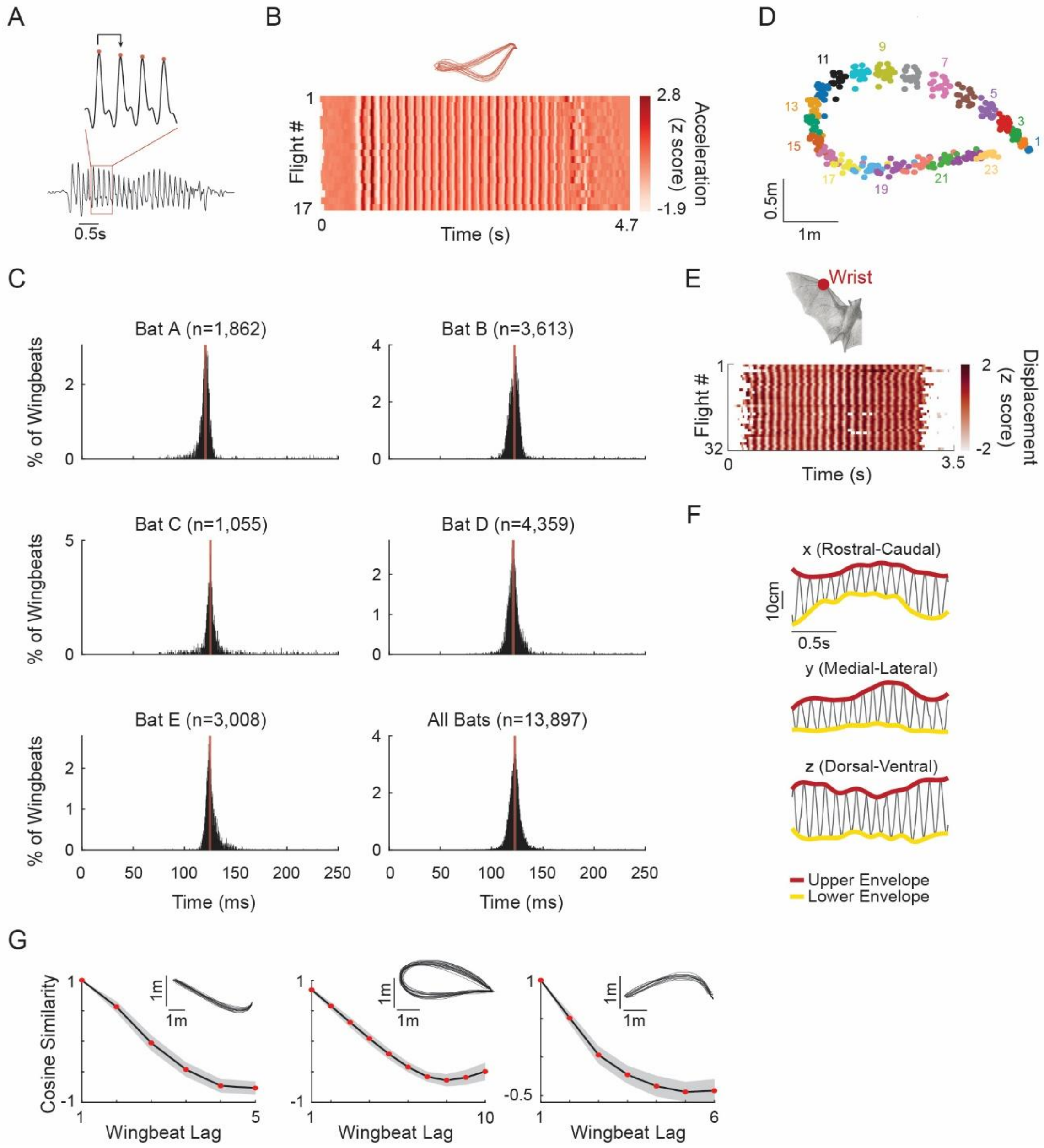


**Figure S4. Wingbeat period variability and adaptation. A)** Wingbeat detection. Bottom: Z-axis acceleration during one example flight. Top: Four wingbeat cycles with detected peaks marked in red. Arrow denotes one wingbeat cycle. **B)** Aligned acceleration traces (z-axis) of all flights on one flight path (z scored). All flights on the same path are shown above. **C)** Distribution of all wingbeat durations detected for each bat and pooled across bats. Number of wingbeats is indicated. **D)** Positional reproducibility across wingbeats. Each dot represents the mean position during a wingbeat (top-down projection), colored by sequential wingbeat number. Only odd numbers are depicted for clarity. Note the tight reproducibility of each wingbeat along the flight trajectory. **E)** Wrist displacement (z scored) in body-centered coordinates across flights of one path. White gaps indicate frames where the key point was not detected. **F)** Example upper (red) and lower (yellow) displacement envelopes for one flight (shown along three axes: rostral-caudal, medial-lateral and dorsoventral axes). **G)** Cosine similarity of adaptation vectors (Figure 1) as a function of wingbeat lag for three example paths (inset). Red: mean; gray: SD.

Figure S5. Two hypothetical computational regimes of motor cortex

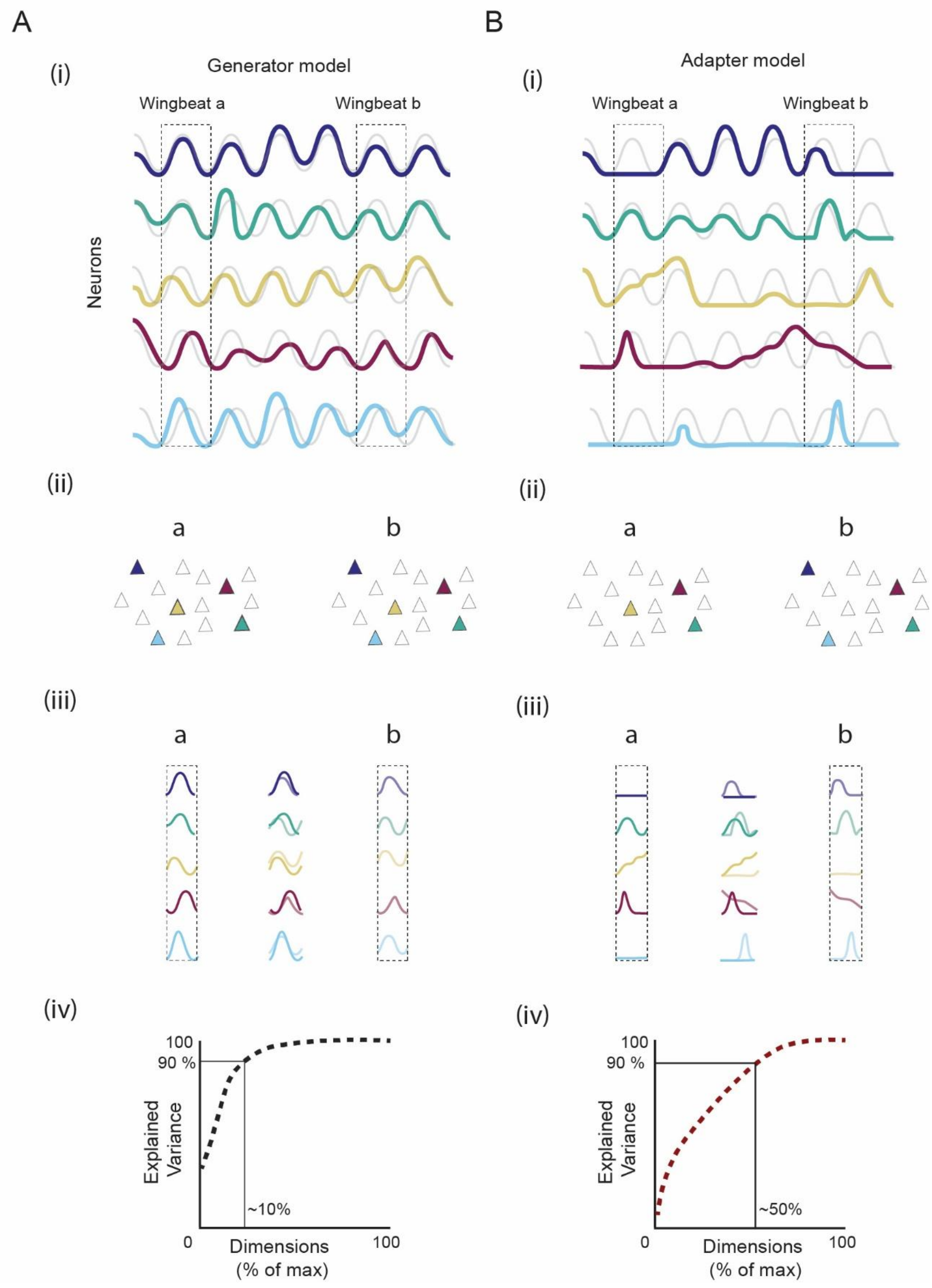

**Figure S5. Two hypothetical computational regimes of motor cortex.** Each wingbeat consists of a shared rhythmic cycle plus cycle-specific adaptations. We consider two regimes for how motor cortex might represent these components. In Regime 1. **A**) neural activity faithfully reflects the full movement: most variance tracks the shared cycle, with smaller firing rate modulations encoding adaptations. In Regime 2. **B**) neural activity disproportionately reflects cycle-specific adaptations and variance is less dominated by the shared rhythm. These two regimes result in distinct predictions of neural patterns from the single neuron to the ensemble activity (i-iv). **(i)** Simulated activity of five hypothetical neurons across six wingbeats (colored traces; gray: underlying wingbeat cycle). In Regime 1, neurons are consistently active on each cycle with firing rate modulations on each wingbeat. In Regime 2, neurons fire more selectively, engaging most selectively when specific adaptations are needed on top of the rhythmic wingbeat cycle. **(ii)** Neuron recruitment for two example wingbeats (a, b). Regime 1 recruits most neurons every cycle; Regime 2 recruits different subsets depending on the adaptations of each wingbeat. **(iii)** Overlay of single-neuron activity across wingbeats 'a' and 'b'. In Regime 1, activity is similar across wingbeats because it is dominated by the shared cycle. In Regime 2, activity differs substantially between wingbeats because it primarily reflects wingbeat-specific demands. **(iv)** These regimes make opposing predictions about population dimensionality (cumulative explained variance from PCA; x-axis normalized to number of neurons). Regime 1 predicts low dimensionality with few dimensions capturing most variance as activity is broadly shared. Regime 2 predicts high dimensionality; more dimensions are needed because different wingbeats recruit distinct neural population patterns.

Figure S6. Wingbeat group clustering and neuronal wingbeat similarity

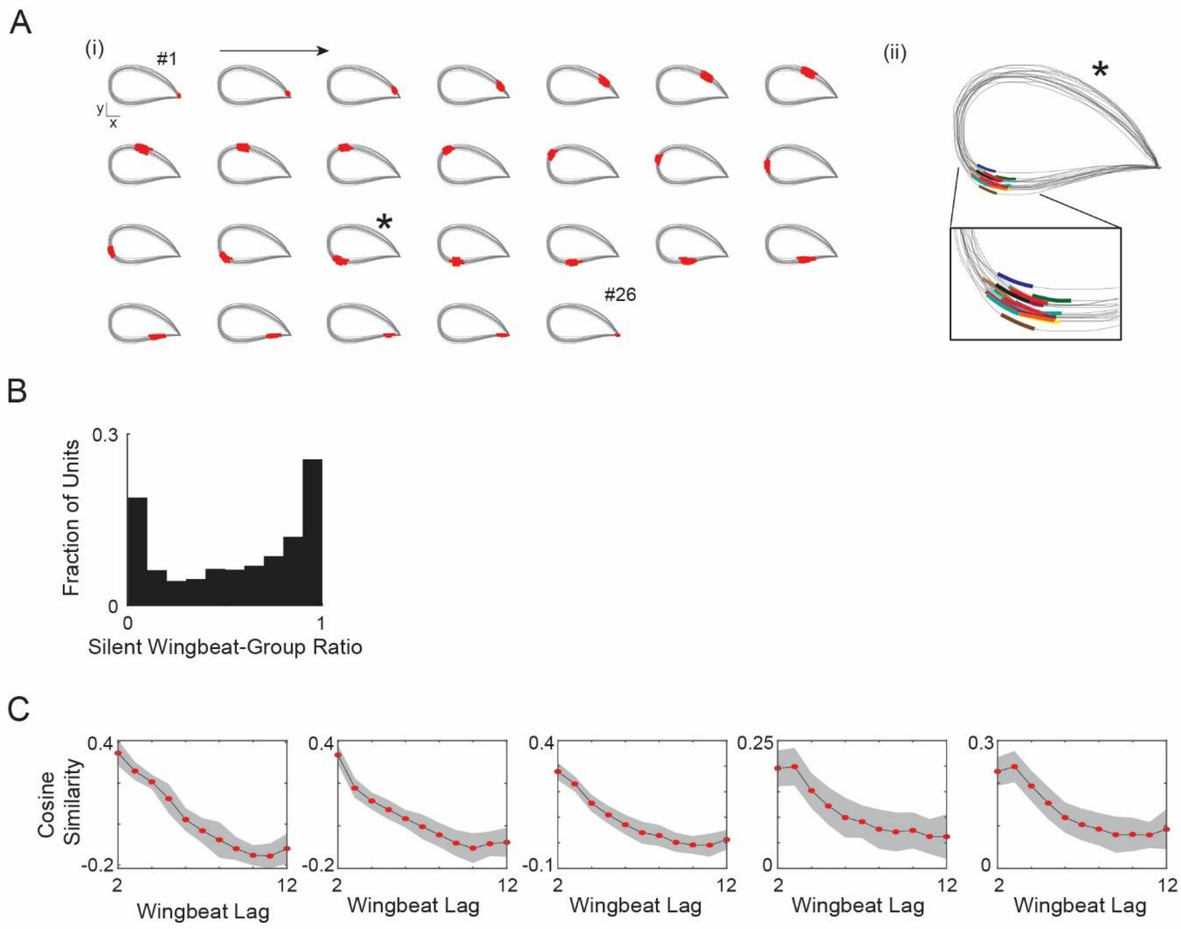


**Figure S6. Wingbeat group clustering and neuronal population wingbeat similarity. A)** (i) Example flight trajectory containing 26 wingbeat groups identified by clustering (Methods). Each panel shows one wingbeat group; gray lines depict the top-down projection of all positions across all flights along this trajectory, and positions occupied during each wingbeat group are indicated in red. Wingbeat groups ordered chronologically (left to right, top to bottom, as indicated by the arrow). Asterisk marks the wingbeat group shown in (ii); first (#1) and last (#26) wingbeat group are indicated. (ii) Top: enlarged view of wingbeat group #17 (asterisk) with individual wingbeats indicated (colors). Bottom: close-up on individual wingbeats. **B)** Distribution of silent wingbeat-group ratios across all flight-modulated units. A wingbeat group was classified as silent for a given unit if the unit emitted zero spikes in ≥ 65% of the wingbeats in that group. N = 1,180 units, p = 17 trajectories, N = 5 bats. **C)** Cosine similarity between the mean-subtracted firing rate vector at each wingbeat and forward-lagged wingbeat vectors, computed across all flights of a given trajectory. Five trajectories with over 100 co-recorded units are shown. Red dots, mean across flights; gray shading, SD across flights.

#### Figure S7. Wingbeat phase decoding generalizes across flight paths

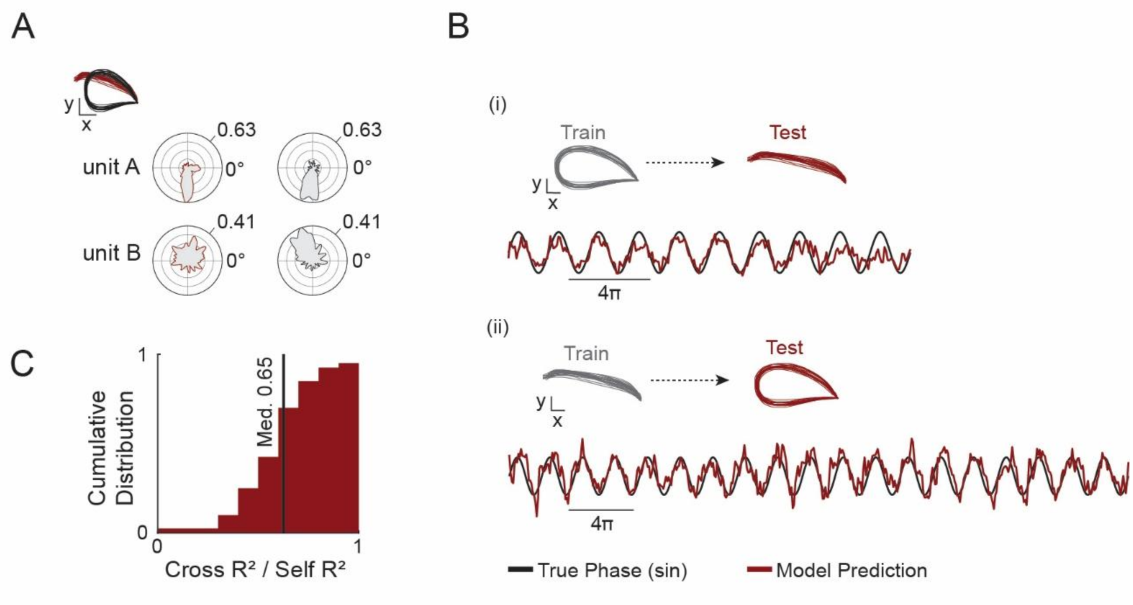


**Figure S7. Wingbeat phase decoding generalizes across flight paths. A)** Polar tuning plots for two example units recorded on two different flight paths (depicted top-left). Red outlines, activity during the red path; black outlines, activity during the black path. Numbers indicate the probability density range (computed jointly across both paths to allow direct comparison). Unit A shows similar phase tuning across both paths, consistent with kinematic invariance. Unit B is tuned only during the black path, indicating sensitivity to path-specific kinematics. **B)** Cross-path wingbeat phase decoding using ordinary least squares regression. Top: schematic indicating train and test path identity. Bottom: cross path predicted sine component of wingbeat phase (red) overlaid on the true sine component (black). (i) Model trained on one path and tested on a kinematically distinct path. (ii) Train and test path pairs swapped. **C)** Cumulative distribution of cross-path $R^2$ normalized to self-$R^2$ (computed using the training set as the test set). Median normalized $R^2 = 0.65$; n = 22 paths; N = 5 bats.

#### Figure S8. PCA lower-bound dimensionality estimates and GPFA summary

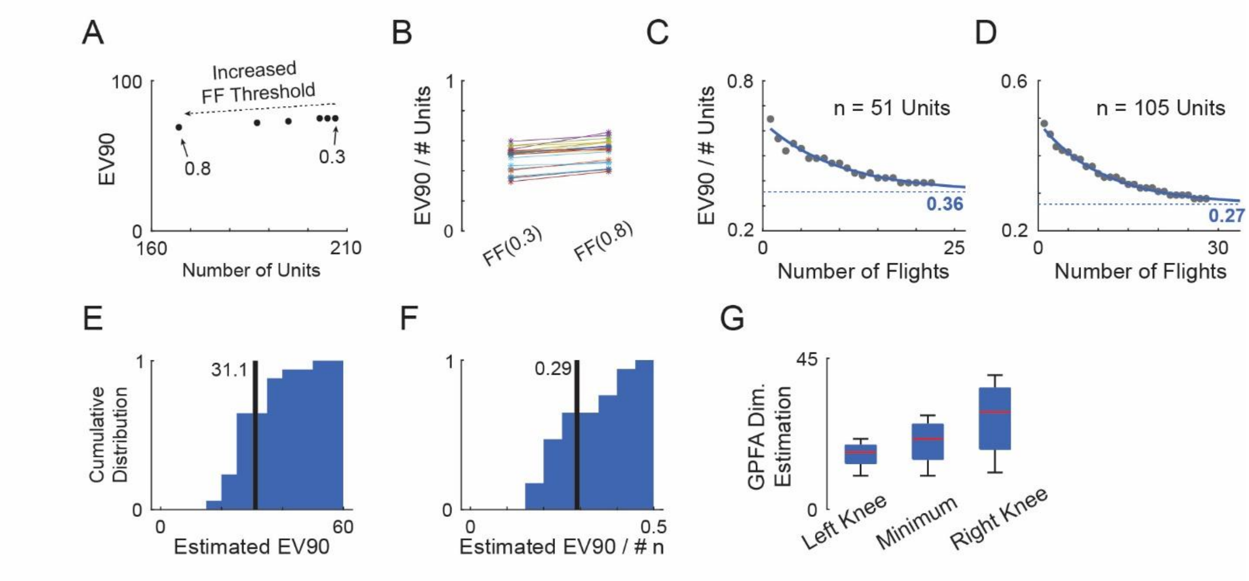


**Figure S8. PCA lower-bound dimensionality estimates and GPFA summary. A)** Effect of unit stability threshold on EV90 (number of PCs required to explain 90% of variance). Each point represents a different firing-rate stability threshold (FFth, ranging from 0.3 to 0.8; Methods), which controls the percentage of wingbeat groups in which a unit must show consistent firing rates to be included. Note that increasing the threshold reduces the number of included units while modestly affecting EV90. This indicates the most consistent units are not highly similar but maintain a high diversity of activity patterns. **B)** Normalized EV90 (EV90/n, where n is the number of units) across all flight paths for FFth = 0.3 and 0.8. Each line represents one path; n = 22 paths. **C–D)** Normalized EV90 as a function of the number of subsampled flights for two example paths (n = 51 and 105 units, respectively). Blue curve fitted exponential decay model; dashed line, asymptotic dimensionality estimate. **E)** Distribution of asymptotic EV90 estimated from the noise model across all paths with a minimum of 50 units. Black line, mean EV90 = 31.1 ± 8.8 (mean ± SD); n = 17 paths. **F)** Same as (E) but for normalized EV90. Black line, mean normalized EV90 = 0.29 ± 0.09 (mean ± SD); n = 17 paths. **G)** Box plots of estimated shared-subspace dimensionality using leave-unit-out cross-validated GPFA. The minimum cross-validated log-likelihood was used to estimate dimensionality, with a 1% threshold above the minimum defining the left and right range (Knees). Mean dimensionality (minimum) = 21; IQR = 10.25; n = 13 paths.

| Bat ID | Session | Total Units | Active Units | Flight Modulated |
|---|---|---|---|---|
| A | 1 | 179 | 144 | 144 |
| A | 2 | 116 | 93 | 90 |
| A | 3 | 99 | 76 | 66 |
| A | 4 | 74 | 59 | 48 |
| B | 5 | 376 | 321 | 269 |
| B | 6 | 205 | 172 | 159 |
| B | 7 | 93 | 79 | 79 |
| B | 8 | 65 | 53 | 51 |
| C | 9 | 197 | 156 | 97 |
| C | 11 | 82 | 63 | 39 |
| D | 12 | 143 | 118 | 116 |
| D | 13 | 110 | 91 | 86 |
| D | 14 | 90 | 67 | 58 |
| E | 15 | 195 | 160 | 155 |
| E | 16 | 128 | 108 | 105 |

**Supplementary Table 1: Unit counts by recording session and functional category.** Each row represents one recording session from one bat (A-E; session number is indicated). **Total Units:** all sorted single units recorded during the session. **Active Units:** active units during flight (>2 spikes in >70% of flights for at least one path). **Flight Modulated Units:** active units showing significant firing rate modulation along at least one path (Methods). Sessions included met minimum thresholds of ≥35 simultaneously recorded active units and ≥8 flights per path. N=16 sessions from 5 bats.
